# The genetic basis of predation resistance in *Pseudomonas* species associated with the bactivorous soil amoeba *Dictyostelium discoideum*

**DOI:** 10.1101/2024.08.09.607352

**Authors:** Margaret I. Steele, Jessica M. Peiser, Simon P. M. Dawson, David C. Queller, Joan E. Strassmann

## Abstract

Predation is likely to influence the function of bacterial communities and the evolution of bacterial pathogens, because characteristics that permit escape from predators often overlap with traits used for biocontrol of plant pathogens, virulence, or even bioremediation. Soil bacteria are preyed upon by a variety of microorganisms, including the amoeba *Dictyostelium discoideum,* which has led some strains to evolve resistance. We identified genes required for three *Pseudomonas* species associated with *D. discoideum* to evade predation by screening more than 6,000 transposon mutants for loss of resistance. One species required a variety of genes including toxins and secondary metabolism genes, but the other two appear to have functionally redundant mechanisms of resistance, since disruption of genes with pleiotropic effects was required to render them susceptible. We determined that GacA, which positively regulates secondary metabolism, is required for resistance in all three species. Predation resistance also appears to be a social trait based on enrichment of cooperative genes in one species and rescue of mutants by wild type in another. Many genes required for resistance are conserved among both resistant and susceptible species, but several are found in few genomes and some of these have homologs in distantly related species. Gain and loss of resistance appears to be a dynamic process in which regulatory and structural genes are well conserved across species, the specific toxins they regulate may be lost in the absence of predators, and new toxins may be acquired through horizontal gene transfer.

## Introduction

Soil ecosystems are home to diverse populations of bacteria that serve numerous ecologically important roles, such as nutrient cycling, bioremediation, pathogenicity, biocontrol of plant pathogens, and mutualisms with plants and other microbes (Thies and Grossman 2023). Among soil bacteria, species belonging to *Pseudomonas* are particularly well studied due to their diversity, relevance to plant and animal health, and genetic tractability (Garrido-Sanz et al. 2016; Lalucat et al. 2020). These species have evolved myriad ways of interacting with other organisms. Several species within the *Pseudomonas fluorescens* group, including *Pseudomonas protegens,* are considered plant growth-promoting rhizobacteria due to their ability to inhibit phytopathogenic bacteria and fungi (Ellis et al. 2000; Raaijmakers et al. 2002; Raaijmakers et al. 2009). *Pseudomonas aeruginosa* lives in soil and water but opportunistically infects humans (Green et al. 1974; Schroth et al. 2018). Other species are pathogens of plants (Xin et al. 2018) and insects (Vodovar et al. 2006). However, these bacteria are also subject to predation by a variety of soil protists and nematodes, necessitating mechanisms of self-defense (Jousset 2012; Martins et al. 2022; Jiang et al. 2023).

Bacteria employ a variety of strategies to defend themselves from predators, including biofilm formation, motility, and secretion of proteins and metabolites (Matz and Kjelleberg 2005). Bacterial characteristics that permit escape from predation often overlap with traits used in biocontrol of plant pathogens, virulence, or even bioremediation (Jousset 2012). Because some mechanisms used to inhibit fungal competitors also affect protists, strains used for biocontrol tend to resist protozoan grazing (Jousset et al. 2006; Song et al. 2015; Amacker et al. 2020).

These mechanisms include secretion of secondary metabolites and exoproteases, which are positively regulated by the GacA-GacS two-component system (Seaton et al. 2013; Stallforth et al. 2013; Amacker et al. 2020; Götze et al. 2023; Pflanze et al. 2023). In many species, toxic proteins can also be injected directly into phagocytic cells using a Type III secretion system (T3SS) (Pukatzki et al. 2002; Basso et al. 2017) or Type VI secretion system (T6SS) (Boak et al. 2022). Furthermore, genes that contribute to the virulence of *P. aeruginosa* in mammalian hosts also confer resistance to predation by amoebae (Pukatzki et al. 2002) and exposure to protists increases the infectivity of *Vibrio cholerae* (Espinoza-Vergara et al. 2020; Hoque et al. 2022), suggesting that predation may drive the evolution of bacterial virulence (Matz and Kjelleberg 2005).

The protist *Dictyostelium discoideum* is a generalist predator that uses phagocytosis to prey on Gram-positive and Gram-negative bacteria from multiple phyla (Brock et al. 2018; Rashidi and Ostrowski 2019; Shreenidhi et al. 2024). *D. discoideum* consumes bacteria as a unicellular amoeba and forms multicellular fruiting bodies in response to starvation. *D. discoideum* also acts as a host for a variety of endosymbionts (DiSalvo et al. 2015; Haselkorn et al. 2021) and pathogens (Taylor-Mulneix et al. 2017; Butler et al. 2020) and has been used to identify virulence factors in human pathogens (Pukatzki et al. 2002; Hasselbring et al. 2011).

Resistance to predation by *D. discoideum* appears to have evolved many times in bacteria (Brock et al. 2018) and in *Pseudomonas* specifically (Steele et al. 2023). Furthermore, predation-resistant *Pseudomonas* sporadically infect *D. discoideum* fruiting bodies and some species promote secondary infections by prey bacteria (Brock et al. 2018; Steele et al. 2023). Both predation-resistant and susceptible *Pseudomonas* encode a variety of secretion systems and secondary metabolite biosynthetic gene clusters, but no one set of such genes is found in all predation-resistant strains or exclusively in predation-resistant strains (Steele et al. 2023).

In this study we screened more than 6,000 transposon mutants to identify genes required for predation-resistance in three *Pseudomonas* species isolated from fruiting bodies of wild *D. discoideum* clones. This analysis allowed us to investigate the extent to which different lineages have converged on similar mechanisms of resistance. The strains used were *Pseudomonas* sp. 20P_3.2_Bac4 (hereafter referred to as P324), *P. protegens* 18P_8.2_Bac1 (P821), and *Pseudomonas* sp. 6D_7.1_Bac1 (P711), which are each more closely related to susceptible bacteria than they are to one another, suggesting they may represent independent evolutions of resistance (Steele et al. 2023). *Pseudomonas* sp. P324 and *P. protegens* P821 both appear to have functionally redundant mechanisms of predation resistance, as only transposon insertions in genes that affect multiple pathways compromised resistance. In contrast, many genes in *Pseudomonas* sp. P711 work together to confer resistance. Genes that contributed to resistance included T6SS effectors, two component systems, transcriptional regulators, and transporters. Furthermore, *gacA*, the global regulator of secondary metabolism, was required for resistance in all three species, suggesting that resistance relies on conserved regulatory machinery, even if the toxins produced vary between species.

## Results and Discussion

### Distribution of predation resistance genes in *Pseudomonas*

To investigate how three different *Pseudomonas* isolates evade predation by *D. discoideum,* we screened more than 2,000 transposon mutants from each species for loss of predation resistance. Each of the three isolates encodes approximately 6,000 genes in its genome. In *P. aeruginosa*, there are fewer than 500 essential genes for which disruption by a transposon is fatal (Poulsen et al. 2019), which means that the 2,000 mutants we screened will not include disruptions of every non-essential gene. The genes we identified therefore represent a subset of the genes required for predation resistance. We identified 5 proteins important for resistance in *Pseudomonas* sp. P324 (Table 2) and 71 proteins in *Pseudomonas* P711 (Table S1). None of the *P. protegens* transposon mutants that we tested were susceptible to predation, suggesting P821 may employ redundant strategies so that disruption of a single gene is unlikely to lead to loss of resistance. Similarly, at least 3 the 5 genes identified in *Pseudomonas* P324 (Table 2) affect the expression or function of many other proteins. These genes are also present in both resistant and susceptible *Pseudomonas* isolates, indicating that they do not directly confer resistance but may regulate the proteins that do (Figure 1). The fact that we did not identify any mutants with transposon insertions in genes that directly contribute to resistance suggests that multiple processes must be disrupted for P324 to become susceptible to predation. In contrast, several of the genes identified in P711 were toxins or located in clusters of secondary metabolite biosynthesis genes, which suggests that multiple mechanisms work together to confer resistance and disruption of any one pathway is enough to render the bacteria susceptible to predation. While many of the genes identified in P711 are similarly found in all the genomes we analyzed, several have more limited distributions (Figure 1). These include genes associated with secondary metabolite biosynthesis, putative toxins and social genes, and genes of unknown function. Furthermore, some of these genes have no homologs in the genomes of close relatives but are present in distantly related *Pseudomonas* species. It is possible these genes were present in the common ancestor and lost in all but these few strains, but for some genes this would require many losses (Figure 1, Figure S1). Two examples of this are a toll/interleukin-1 receptor domain-containing protein that shares 96.6% amino acid identity with a protein from *P. brenneri* and a MBL fold metallo-hydrolase protein that shares 79.6% amino acid identity with a protein encoded by *P. panipatensis.* Additionally, phylogenies of four predation resistance genes are not congruent with a phylogeny built from 78 conserved core proteins, indicating that their evolutionary history differs from that of the rest of the genome (Figure S2). This result suggests horizontal gene transfer may be important for the acquisition of a predation resistant phenotype.

**Figure 1.**
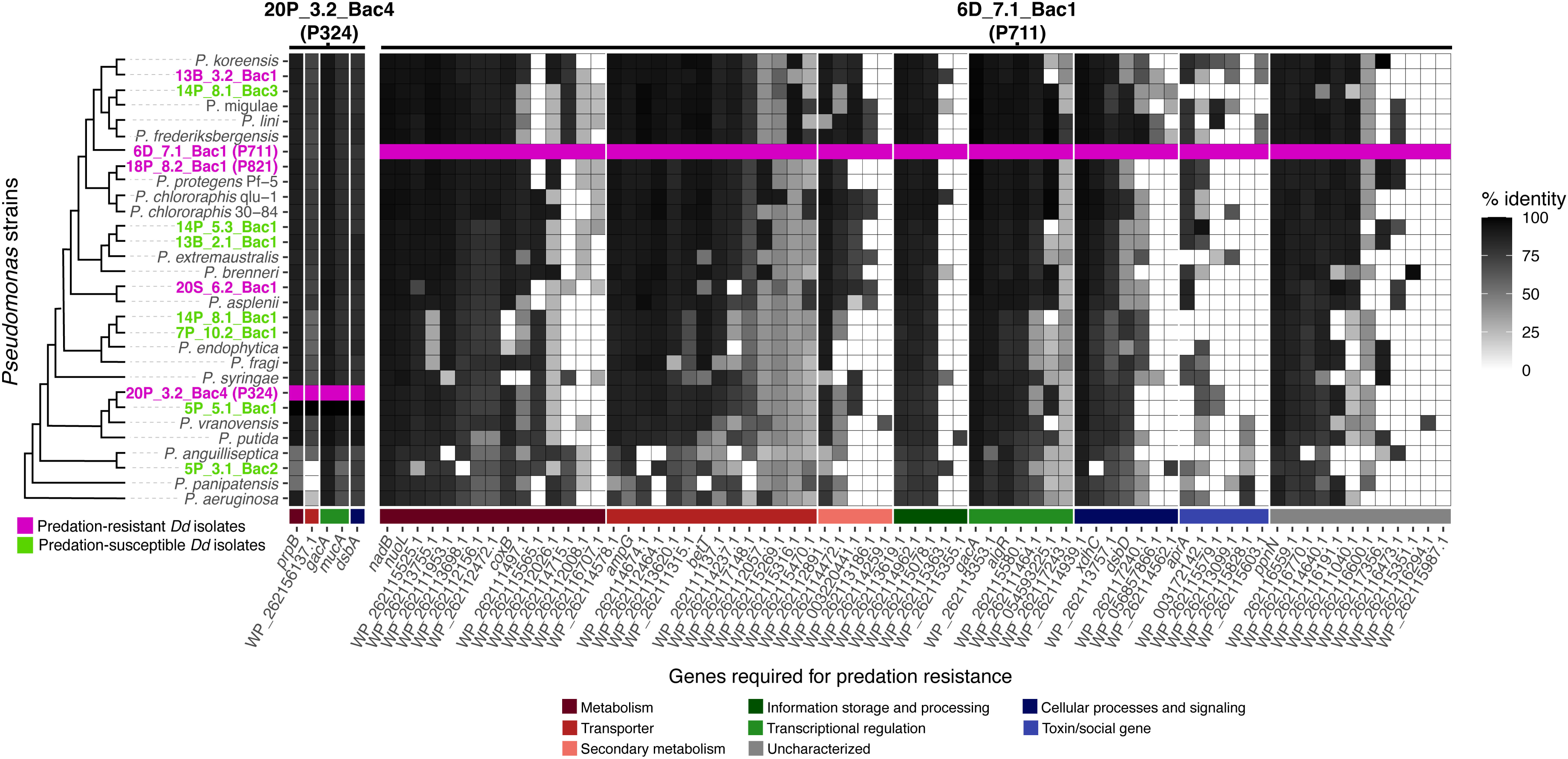
Many genes required for predation resistance are conserved across resistant and susceptible strains, but some have more limited distributions. BLASTp was used to search *Pseudomonas* genomes for homologs to proteins identified as required for predation resistance in P324 or P711. A genome phylogeny built from 1703 gene trees is shown on the left. Names of strains isolated from *D. discoideum* are colored, with predation-resistant strains in magenta and predation-susceptible strains in green. The x-axis shows the names of proteins identified by screening transposon mutants for loss of predation resistance. Proteins are grouped according to strain of origin (P324 or P711) and predicted function, shown by the colored bar below the heat map, then sorted based on the average nucleotide identity of BLAST hits across all genomes so that the most conserved genes are on the left. The row highlighted in magenta contains 100% identity matches between predation resistance genes and their genome of origin.

**Table 1.**
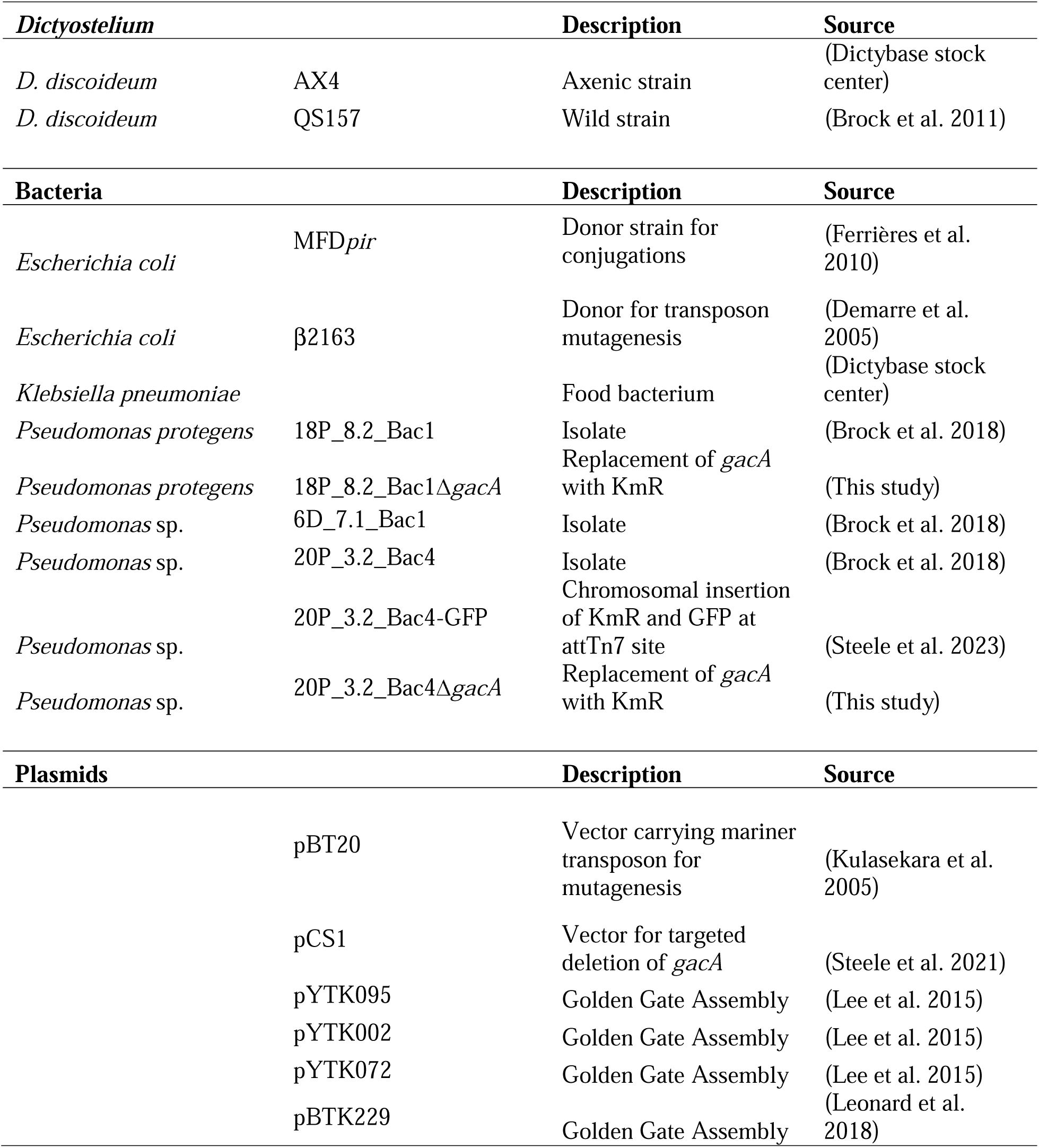
Strains and plasmids used in this study.

**Table 2.** *Pseudomonas* P324 transposon mutants with increased susceptibility to predation by *D. discoideum*.

| **Mutant ID** | **RefSeq Protein** | **Annotation** | **Description (emapper)** | **COG category** | **Protein Localization (pSORTb)** |
| --- | --- | --- | --- | --- | --- |
| P324 1-B12 | WP_095159042.1 | *dsbA* | Thiol disulfide interchange protein | O | Periplasmic |
| P324 16-D2 | WP_262154216.1 | *prpC* | Belongs to the citrate synthase family | C | Cytoplasmic |
| P324 21-E7 | WP_262154406.1 | *rseA/mucA* | Anti sigma-E protein<br>RseA | T | Cytoplasmic<br>Membrane |
| P324 3-D5 | WP_262156137.1 | ABC transporter substrate-binding protein | Amino acid ABC transporter substrate-binding protein | ET | Cytoplasmic |
| P324 6-C11 | WP_095159785.1 | *gacA/uvrY* | Two component transcriptional regulator, LuxR family | K | Cytoplasmic |
| P324 7-A10 | WP_262154406.1 | *rseA/mucA* | Anti sigma-E protein<br>RseA | T | Cytoplasmic<br>Membrane |

The predation resistance genes that we identified are functionally diverse. A COG functional enrichment analysis revealed that no categories were significantly over or underrepresented among the genes identified in our screen (Figure 2). This may be due to the small number of genes in the analysis, but it also suggests that many different functions are required for *Pseudomonas* to escape predation by *D. discoideum*. We examined the predicted subcellular locations of proteins identified in our screen using pSORTb (Yu et al. 2010) and found that the proportion of proteins assigned to each location was similar to the genome as a whole, indicating that predation resistance genes are not more likely than expected to encode secreted proteins (Figure 3A). This emphasizes that predation resistance depends on a network of proteins, including structural, regulatory, and metabolic genes.

**Figure 2.**
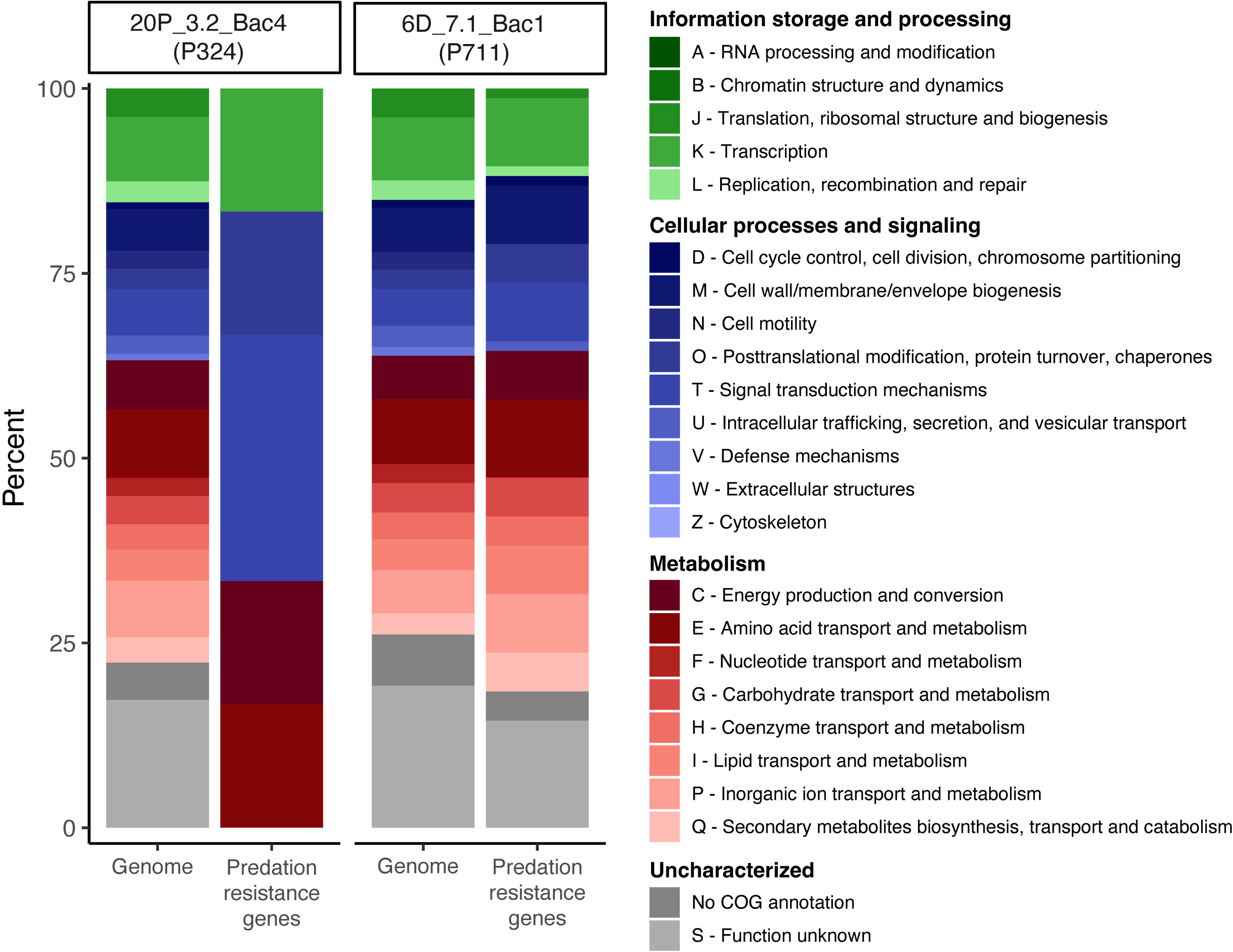
Predation resistance genes in *Pseudomonas* P324 and P711 belong to diverse functional categories. eggNOG-mapper was used to assign COG functions to all genes in the genome and genes identified as required for predation resistance. Fisher’s Exact Test was used to test for significant differences in the proportion of genes belonging to each category.

**Figure 3.**
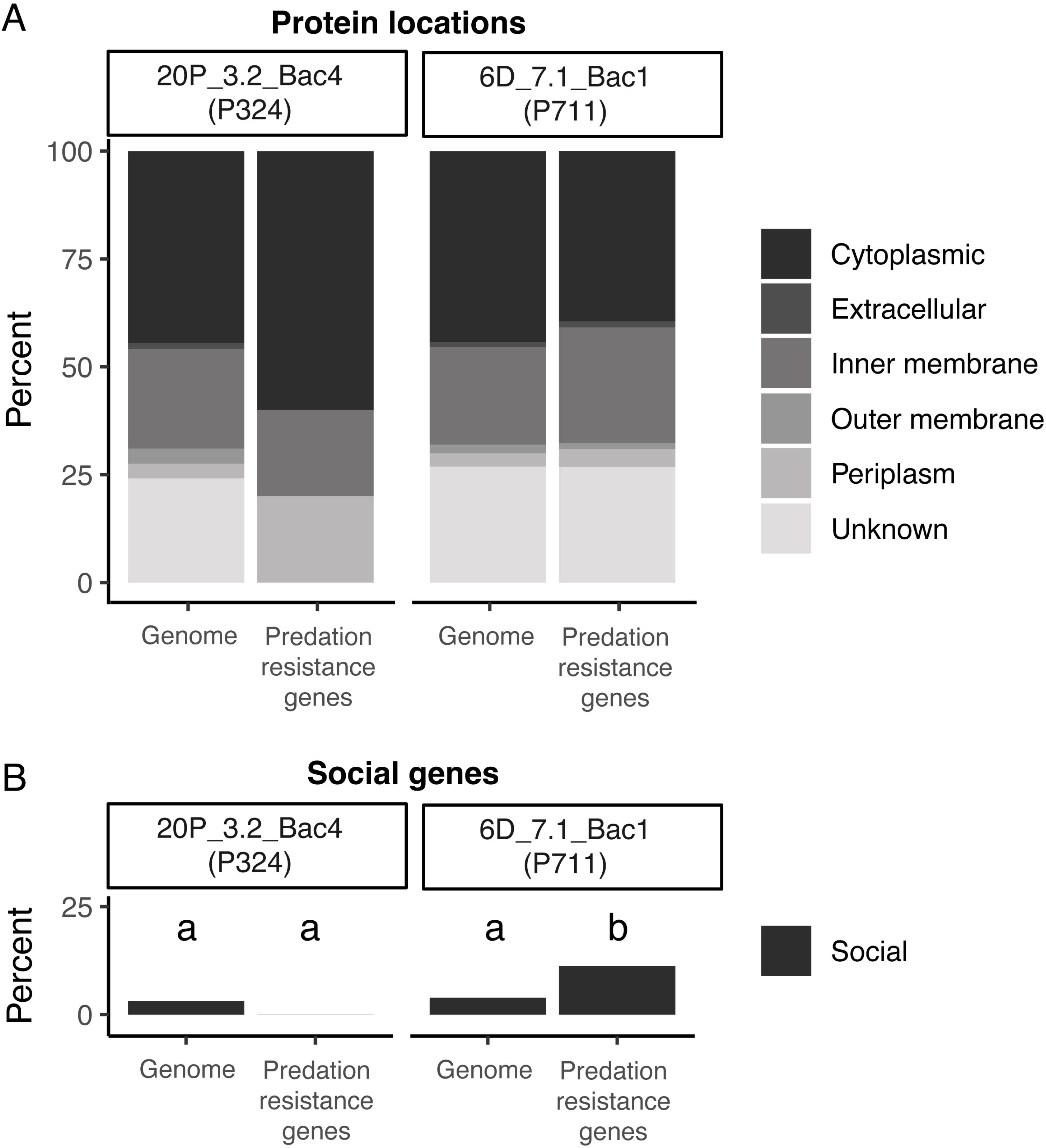
Social genes are overrepresented among predation resistance genes in *Pseudomonas* P711. (A) Cellular locations of proteins were predicted using pSORTb. (B) SOCfinder was used to predict genes with social functions. Fisher’s Exact Test, FDR correction for multiple comparisons, q=0.05.

Our three predation resistant *Pseudomonas* species inhibit consumption of prey bacteria in coculture (Steele et al., 2023), potentially due to the secretion of toxic proteins or metabolites, which has been observed in other *Pseudomonas* species (Raaijmakers et al. 2009; Jousset 2012). To determine whether the predation resistance genes that we identified are likely to benefit other members of the community, we used SOCfinder to annotate cooperative genes. SOCfinder conservatively identifies such genes based on sequence motifs associated with extracellular localization, secondary metabolite biosynthetic gene clusters, and homology to known social genes (Belcher et al. 2023). We found that social genes are significantly enriched among predation resistance genes in P711 (Figure 3B). This may be an underestimate, because several toxins and secondary metabolite biosynthetic genes were not identified as cooperative but may benefit nearby bacteria. The lack of significant enrichment of social genes in PP324 is likely because the small number of genes that were identified indirectly affect cooperative traits. For example, *gacA*, which positively regulates secondary metabolism, and *dsbA*, which is required for the function of many secreted proteins, are not annotated as cooperative genes. Furthermore, we found that of three susceptible P324 mutants that we tested, two were partially rescued from predation in coculture with WT, supporting the conclusion that predation resistance is at least partially a social trait (Figure 4).

**Figure 4.**
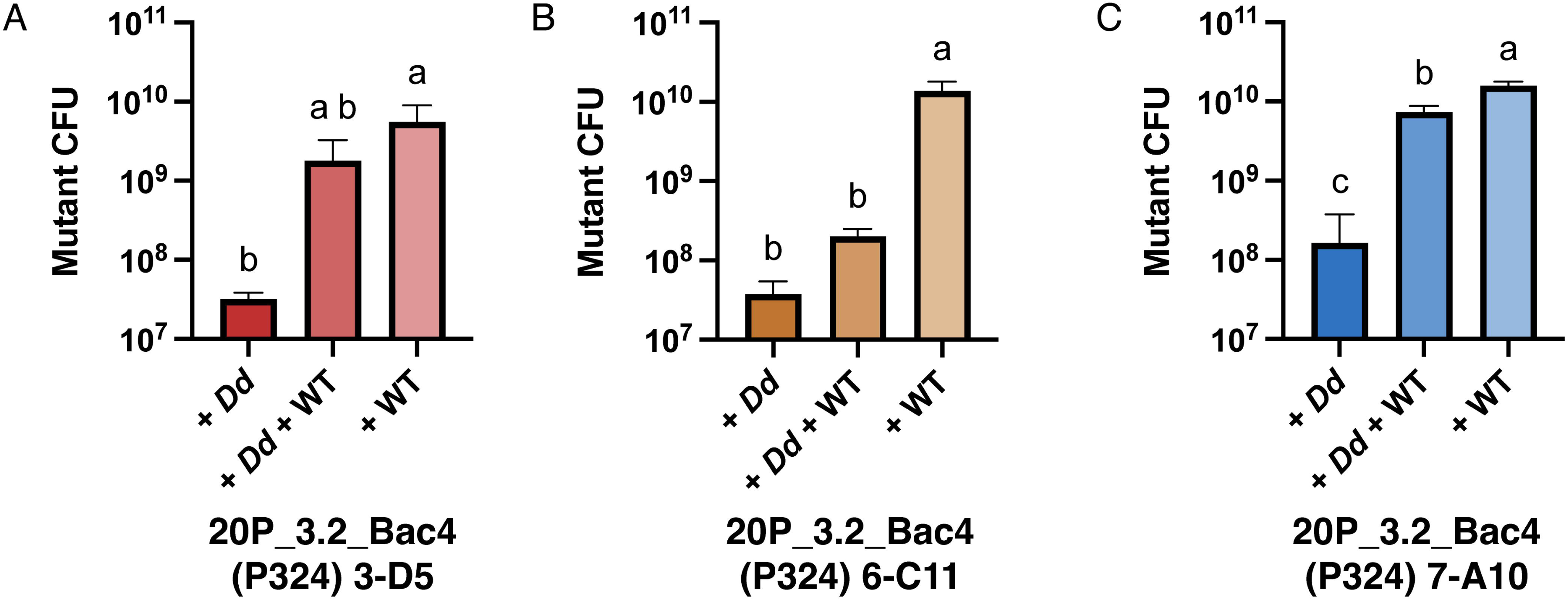
Coculture with wild type partially rescues some *Pseudomonas* P324 mutants from predation by *D. discoideum.* *Pseudomonas* P324 mutants were grown with *D. discoideum* QS157 (+*Dd*), QS157 and wild type P324 (+*Dd* +WT), or with wild type alone (+WT). (A) 3-D5, which has a transposon insertion in an ABC transporter substrate-binding protein, (B) 6-C11, which has a transposon insertion in *gacA*, and (C) 7-A10, which has a transposon insertion in *rseA.* One-way ANOVA with Tukey’s multiple comparison test, p<0.05.

### Secondary metabolism contributes to resistance in all three species

Secondary metabolism plays a significant role in the ability of all three *Pseudomonas* strains to evade predation by *D. discoideum.* Among the susceptible mutants, we identified transposon insertions in the *gacA* gene in both P324 and P711 (Table 2, 3, S1). GacA is the response regulator in the GacA-GacS two component system, which regulates the expression of hundreds of genes, including positive regulation of secondary metabolism (Hassan et al. 2010). A nonsense mutation in *gacA* has previously been shown to render *P. protegens* Pf2 susceptible to predation by *D. discoideum* (Stallforth et al. 2013; Inglis et al. 2018) and deletion of *gacS* increases consumption of *P. fluorescens* by a variety of protists and nematodes (Jousset et al. 2006; Jousset et al. 2009; Burlinson et al. 2013). *Pseudomonas* sp. P324 and *P. protegens* P821 *gacA* mutants also grew to higher abundance than wild type in the absence of predation (Figure S3). This is consistent with reduced metabolic load and increased growth rate reported for *gacA* mutants in other *Pseudomonas* species and demonstrates that loss of *gacA* can be beneficial to fitness in the absence of predators or competitors (Driscoll et al. 2011; Yan et al. 2018).

Further emphasizing the importance of secondary metabolism, in P711, we identified seven susceptible mutants with transposon insertions in five of the thirteen secondary metabolite biosynthesis gene clusters predicted by antiSMASH (Table 3) (Medema et al. 2011; Blin et al. 2021). Four of the susceptible mutants had transposon insertions in genes in non-ribosomal peptide synthetase (NRPS) clusters. We identified two genes important for predation resistance within a NRPS/betalactone protocluster, which includes genes with similarity to 100% of genes in the pseudomonine biosynthesis cluster. Pseudomonine synthase produces multiple siderophores and siderophore-like molecules based on substrate availability (Wuest et al. 2009). Other *Pseudomonas* siderophores inhibit fungal growth by sequestering iron (Kloepper et al. 1980; DÉFAGO and HAAS 1990). *D. discoideum* also requires iron for growth and iron transport is important for resistance to pathogenic bacteria (Peracino et al. 2006; Bozzaro et al. 2013; Buracco et al. 2018), suggesting that siderophore production could be a strategy for inhibiting predator growth. Our screen also found two different genes within a NRPS-like protocluster in which three genes showed similarity to the fragin biosynthetic gene cluster. Fragin has antifungal, antibacterial, and antitumor activity (Murayama et al. 1969).

**Table 3.** *Pseudomonas* P711 transposon insertions in secondary metabolism genes.

| **Mutant** | **RefSeq Protein** | **antiSMASH cluster** | **Annotation** | **COG category** |
| --- | --- | --- | --- | --- |
| P711 8-G5 | WP_019690936.1 | N/A | GacA | K |
| P711 8-H4 | WP_262115525.1 | Hserlactone/<br>butyrolactone | Acetyl-CoA C-acetyltransferase,<br>FacI | I |
| P711 17-D1 | WP_262113186.1 | NRPS-like | Ig-like domain-containing protein | U |
| P711 1-E6 | WP_003220441.1 | NRPS-like | Diiron oxygenase | S |
| P711 14-C6 | WP_262115269.1 | NRPS/betalactone | Efflux RND transporter periplasmic<br>adaptor subunit | M |
| P711 3-E5 | * | NRPS/betalactone | * | * |
| P711 7-H2 | WP_262112891.1 | Redox-cofactor | LysR family transcriptional regulator | K |
| P711 16-C9 | WP_262114472.1 | RiPP-like | DUF692 domain-containing protein | S |
| P711 16-E1 | WP_262114259.1 | T1PKS | Lipoteichoic acid synthase family<br>protein | M |

\* Transposon is located within gene cluster, but the disrupted gene could not be identified

Other secondary metabolite biosynthetic genes that appear to be required for predation resistance in P711 were located within redox-cofactor, a Type I polyketide synthase (T1PKS), and a ribosomally synthesized and post-translationally modified peptide product (RiPP) clusters. One of these genes was a LysR family transcriptional regulator in a redox-cofactor cluster with limited similarity to the biosynthetic cluster for lankacidin C. Lankacidins are non-ribosomal peptides that inhibit protein synthesis in bacteria and interfere with microtubule growth in tumor cells (Ayoub et al. 2016; Cai et al. 2021). Microtubules are required for phagosome maturation in *D. discoideum* (Clarke and Maddera, 2006), which suggests a possible mechanism of action if this biosynthetic cluster encodes a lankacidin-like metabolite. Another mutant had a transposon insertion in a lipoteichoic acid synthetase family protein in a T1PKS cluster. Polyketide synthases are known to produce a variety of products, including phytotoxin, antibiotic, antifungal, insecticide, and antitumor compounds (Bender et al. 1999; Wang et al. 2020). A transposon insertion in an RiPP biosynthetic gene cluster was located in gene encoding a DUF692 domain-containing protein. The product of this gene cluster is unknown, but it is identical to a cluster found in *Pseudomonas brassicacearum*, a plant pathogen (Yang et al. 2020). There was also a mutant with a transposon insertion in a metallo-beta-lactamase superfamily gene that was not annotated by antiSMASH but might be involved in secondary metabolism.

To verify the importance of *gacA* and its downstream genes for predation resistance in our three *Pseudomonas* strains, we deleted the *gacA* gene in P324 and *P. protegens* P821. We quantified loss of predation resistance in the P324 and P821 *gacA* deletion mutants, the P324 and P711 *gacA* transposon mutants, the P711 *aprA* mutant (an exoprotease regulated by GacA), and six mutants with transposon insertions in secondary metabolite biosynthetic gene clusters (Fig. 5). To quantify loss of predation resistance, we normalized the number of bacteria recovered from coculture with *D. discoideum* AX4 to the number of bacteria recovered from monoculture after 5 days. For all three strains, the wild type was fully predation resistant, as we recovered as many bacteria from coculture as from monoculture. Each of the mutants was significantly more susceptible to predation. Similar trends were observed when the same strains were grown with the wild clone *D. discoideum* QS157, which was isolated from soil and has had little opportunity to accumulate adaptations to the laboratory environment (Figure S4).

**Figure 5.**
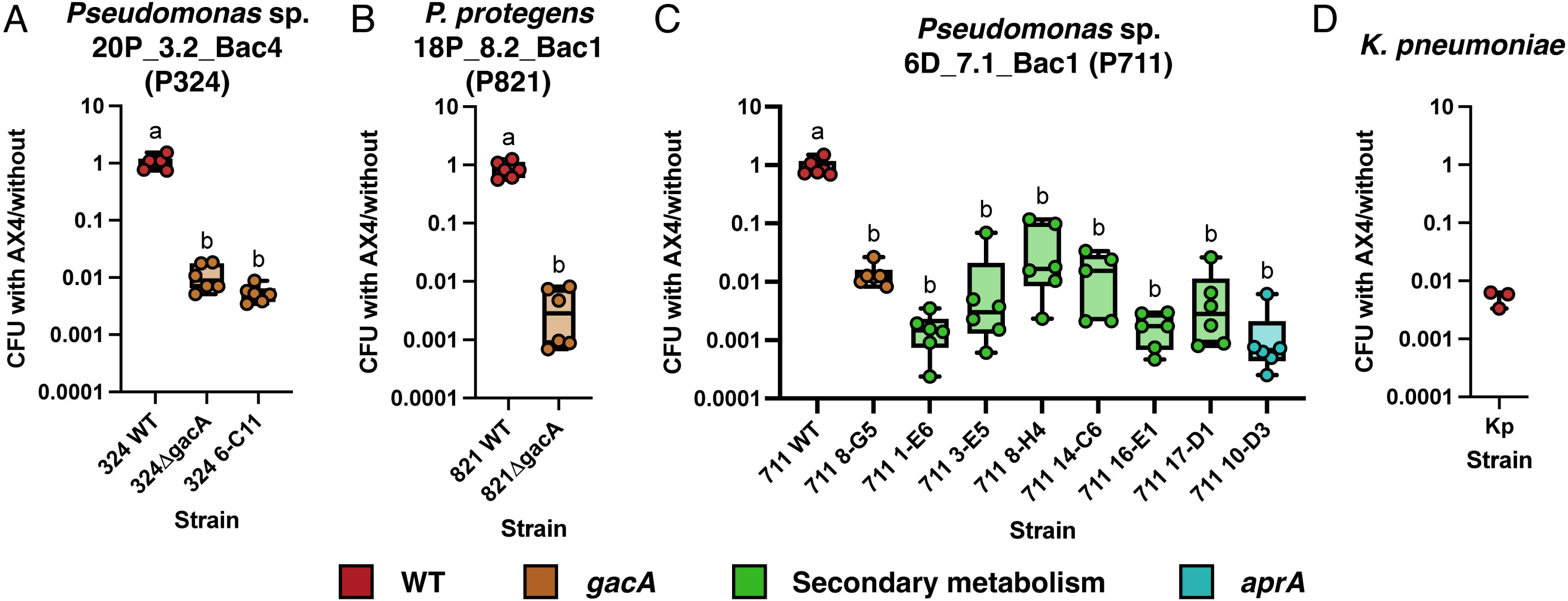
Mutations in *gacA*, secondary metabolite biosynthetic genes, and exoprotease *arpA* make otherwise predation-resistant *Pseudomonas* strains susceptible to predation by *D. discoideum* AX4. (A) *Pseudomonas* P324, (B) *P. protegens* P821, (C) *Pseudomonas* P711, and (D) *K. pneumoniae* (Kp) strains were grown with and without AX4 amoebae on SM/5 agar for 5 d. CFU recovered from coculture with AX4 were normalized by dividing by CFU recovered from monoculture. P324: P324 6-C11, transposon insertion in *gacA*. P711: P711 8-G5, transposon insertion in *gacA*; P711 1-E6, NRPS-like cluster; P711 3-E5, NRPS/betalactone cluster; P711 8-H4, hserlactone/butyolactone cluster; P711 14-C6, NRPS/betalactone cluster; P711 16-E1, T1PKS cluster; P711 17-D1, NRPS-like cluster; P711 10-D3, *aprA*.

### Protein toxins are also required for resistance

Our screen identified multiple toxins that contribute to predation resistance in *Pseudomonas* P711. The best understood is AprA, a secreted serralysin family metalloprotease, which is positively regulated by *gacA* (Hassan et al. 2010) and is toxic against a variety of protists (Jousset et al. 2006; Song et al. 2015; Amacker et al. 2020). A predicted serine protease was also required for resistance. Another putative toxin contained both YopT-type cystine protease and TcdA/TcdB pore forming domains. YopT is a T3SS-associated cysteine protease found in pathogenic *Yersenia* and YopT-like protein folds are found in toxins produced by other bacterial pathogens (Grabowski et al. 2017). TcdA and TcdB are *Clostridium difficile* virulence factors that form pores in membranes to enable toxin delivery (Pruitt et al. 2010). A TcdA/TcdB pore forming domain is found in *fitD*, part of the cluster that encodes the insecticidal toxin Fit in *P. protegens* and *P. chlororaphis* (Ruffner et al. 2015).

Two proteins in clusters of Type VI secretion system (T6SS) effectors were identified as important for predation resistance in *Pseudomonas* P711. T6SS are protein complexes used to inject protein effectors (toxins) through the membranes of prokaryotic or eukaryotic cells. T6SS allow *P. chlororaphis* 30-84 to resist predation by *D. discoideum* and other bactivorous eukaryotes (Boak et al., 2022). One of the genes we found is a hypothetical gene immediately upstream of *hcp,* which forms the needle-like structure ejected by the T6SS. The other is a lysozyme-like protein immediately downstream of *vgrG* and other structural components of the T6SS. We identified two predation-susceptible mutants with different transposon insertion sites in the lysozyme-like effector, confirming the importance of this gene for predation resistance in *Pseudomonas* P711. Both predation-resistant and susceptible *Pseudomonas* strains isolated from *D. discoideum* encode one or more copies of the T6SS structural genes, indicating specific effectors and not the T6SSs themselves are the cause of predation resistance (Steele et al., 2023). Furthermore, T6SS genes in *Pseudomonas fluorescens* Pf-5 are positively regulated by GacA, suggesting loss of T6SS activity may also contribute to the susceptibility of the *gacA* mutants (Hassan et al., 2010).

Our screen also identified homologs to *dsbA* and *dsbD,* which are involved in formation of disulfide bonds in periplasmic proteins, indicating that secreted proteins are required for predation resistance. DsbA, which was required for resistance in P324, catalyzes formation of disulfide bonds in hundreds of periplasmic proteins (Bardwell et al. 1991). Deletion of *dsbA* has pleotropic effects on motility and secretion of virulence factors in *Burkholderia pseudomallei* (Ireland et al. 2014) and in *P. aeruginosa* (Urban et al. 2001; Ha et al. 2003; Heras et al. 2009). DsbD–required in P711–is an inner membrane protein that recycles DsbC, which isomerizes disulfide bonds that were incorrectly oxidized by DsbA (Denoncin and Collet 2013). Inactivation of *dsbA* or *dsbD* is likely to have far reaching effects on the function of periplasmic and extracellular proteins, including proteins that target predators like *D. discoideum*.

### Transporters are required for nutrient uptake and provide information about the environment

In addition to supplying cells with carbon and nitrogen needed for growth, transporters can influence virulence, secondary metabolite biosynthesis, and biofilm formation, all of which are likely to be important for predation resistance. Twelve genes required for predation resistance in our *Pseudomonas* isolates are transporters, including 8 amino acid transporters, 1 carbohydrate transporter, and 3 inorganic ion transporters. Transported ions and molecules are often required for growth, but transporters can also affect gene expression through accumulation of molecules in the cytoplasm or by interacting with two-component systems (Zeng and Charkowski 2021). ATP-binding cassette (ABC) transporters contribute to the virulence of numerous pathogenic bacteria (Lewis et al. 2012; Tanaka et al. 2018).

In P711, we identified two predation-susceptible mutants with transposon insertions in a gene homologous to the putrescine-binding polyamine transporter *potF* and one with an insertion in an ABC transporter permease belonging to the *potC* superfamily. These genes may contribute to predation resistance in multiple ways. For example, putrescine affects virulence and motility in *Proteus mirabilis* (Kurihara(栗原新) et al. 2013) and plant pathogen *Dicketa zeae* (Shi et al. 2019), biofilm formation in environmental bacterium *Shewanella oneidensis* (Ding et al. 2014) and human pathogen *Yersenia pestis* (Patel et al. 2006), and virulence in plant pathogen *Agrobacterium tumefaciens* (Matthysse et al. 1996). In *P. aeruginosa*, polyamines affect antibiotic resistance (Kwon and Lu 2006) and the polyamine spermidine effects expression of T3SS genes (Wu et al. 2012). Furthermore, polyamines may play a role in predator-prey interactions, as *P. fluorescens* SS101 upregulates putrescine biosynthesis in response to grazing by the protist *Naegleria americana* (Song et al., 2015). Additionally, the arginine-binding protein ArtJ, part of the arginine ABC transporter, was also required for resistance in P711. The amino acid arginine is a precursor to polyamines, including putrescine. Arginine is a carbon and nitrogen source for bacteria, but it also promotes biofilm formation in *P. aeruginosa* (Bernier et al. 2011) and *P. putida* (Barrientos-Moreno et al. 2020) and regulates T3SS expression and virulence in enteric pathogens *E. coli* and *Citrobacter rodentium* (Menezes-Garcia et al. 2020).

A number of amino acid transporters not related to polyamines were also required for P711 to evade predation. One of these was a phenylalanine-specific permease. In *P. aeruginosa*, phenylalanine acts as a carbon source and induces biosynthesis of *Pseudomonas* quinolone signal (PQS), a quorum sensing signal required for secretion of virulence factors (Palmer et al. 2010). Transport of phenylalanine increases the availability of chorismite, which is a precursor for biosynthesis of phenylalanine, PQS, and aromatic secondary metabolites (Sterritt et al. 2018). Other amino acid transporters important for resistance include a Carboxylate/Amino Acid/Amine Transporter, a Major Facilitator Superfamily transporter, and two uncharacterized transporters.

The carbohydrate transporter identified in P711 is an AmpG family muropeptide transporter, which likely participates in peptidoglycan recycling. Muropeptides are produced through peptidoglycan breakdown, which occurs during bacterial growth and division, but can also be caused by exposure to lysozymes or antibiotics (Irazoki et al. 2019). *D. discoideum* uses bacteriolytic enzymes to digest bacteria within the phagosome but may also secrete them to kill extracellular bacteria (Ayadi et al. 2024), so the ability to sense damage to the cell wall may be useful for predation resistance.

Another gene required for resistance was homologous to BetT, a choline uptake protein. Bacteria used for biocontrol can slow the growth of fungal plant pathogens by competing with them for choline (Schisler et al. 2006), which can be used as a carbon and nitrogen source, but is also a precursor for the osmoprotectant glycine betaine (Wargo 2013) and induces increased virulence factor activity in *P. aeruginosa* (Lisa et al. 1994; Fitzsimmons et al. 2012). Other ion transporters identified in our screen were an oxalate:formate antiporter and a homolog to the potassium-efflux protein KefC.

In P324, the transporter important for predation resistance is the substrate-binding domain of an amino acid transporter belonging to the HisJ superfamily. This protein may play a role in resistance and infection, as pathogenic *P. syringae* upregulates expression of *hisJ* upon exposure to its host, the bactivorous nematode *C. elegans* (Ali et al. 2022). However, the gene disrupted in P324 is adjacent to an extracytoplasmic-function sigma factor, suggesting that polar effects, in which the transposon affects expression of nearby genes, could contribute to the loss of resistance in this mutant. This would be more consistent with our observation, discussed further below, that the other genes required for resistance in this strain all have pleiotropic effects.

### Responses to the environment are controlled by transcriptional regulators

To evade predation, bacteria must be able to detect changes in the environment and respond through rapid changes in gene expression. As might be expected, among the genes required for predation resistance, 2 genes in P324 and 9 in P711 are involved in regulation of transcription. Two-component systems, comprised of a sensor kinase and response regulator, are one of the mechanisms used by bacteria to respond to environmental signals. Response regulator *gacA*, discussed above, is part of the GacA-GacS two-component system and was required for predation resistance in all three *Pseudomonas* species. Other two-component systems identified in P711 regulate nutrient uptake but also affect expression of genes associated with virulence and predation resistance. This includes *tctE*, the histidine sensor kinase of the TctD-TctE two-component system, which regulates citrate uptake in *P. aeruginosa* and affects biofilm formation, antibiotic resistance, and growth (Taylor et al. 2019). Citrate uptake also increases quorum sensing and pyocyanin biosynthesis in *P. aeruginosa* (Mould et al. 2024). Another gene required for resistance was homologous to *aauR*, the response regulator of the AauR-AauS two-component system, which regulates amino acid uptake and metabolism. In the plant pathogen *P. syringae*, AauR and AauS upregulate expression of T3SS genes in response to host produced aspartate and glutamate (Yan et al. 2020). In *P. aeruginosa,* AauR-AauS contributes to biofilm formation, swarming motility, and virulence (Sultan et al. 2023). We also identified a homolog to GcbA, a PleD-family response regulator. PleD, part of the PleC-PleD two-component system, is a diguanylate cyclase that synthesizes the secondary messenger c-di-GMP in *Caulobacter crescentus* (Paul et al. 2004). In *Pseudomonas,* GbcA promotes attachment to surfaces and reduced motility (Petrova et al. 2014) and c-di-GMP levels affect biofilm formation, with lower levels favoring a mucoid phenotype in *P. fluorescens* (Kessler and Kim 2022) and higher levels promoting biofilm formation in *P. aeruginosa* (Irie et al. 2012).

Not all the transcriptional regulators we identified belonged to two-component systems. Two LysR family transcriptional regulators, which are particularly common in *Pseudomonas* (Reen et al. 2013), also played a role in resistance. One of these was BetR, a transcriptional regulator that controls expression of the choline-O-sulfate utilization gene *betC* (Jovcic et al. 2011). Choline uptake, as described above, was also important for resistance. The other LysR family regulator was located within a predicted redox-cofactor protocluster and may regulate biosynthesis of the secondary metabolite. The other putative transcriptional regulators are diverse, but not as well characterized. Two regulators, a pyridoxal phosphate-dependent aminotransferase and an OmpR family response regulator were each located near another gene that was required for resistance. The last regulator belongs to the xenobiotic response element family of transcriptional regulators.

Two regulatory genes, *mucA* and *algR*, that were required for resistance in P324 and P711 appear to have contradictory effects, likely because these two species prioritize different strategies for evading predation. AlgR, required for resistance in P711, regulates production of the exopolysaccharide alginate, which is involved in biofilm formation and helps protect *P. aeruginosa* from phagocytosis by macrophages and predation by *D. discoideum* (Simpson et al. 1988; Govan and Deretic 1996; Bradbury et al. 2011). Mutations in *algR* result in reductions in alginate biosynthesis, twitching motility, and virulence (Lizewski et al. 2002). Interestingly, two mutants with transposon insertions in *mucA,* a RseA-family anti-sigma factor, were identified in P324. In *E. coli*, RseA sequesters the alternative sigma factor RpoE, which regulates the response to envelope stress (De Las Peñas et al. 1997; Alba and Gross 2004). RseA and RpoE are homologous to anti-sigma factor MucA and sigma factor AlgU in *P. aeruginosa* (Yu et al. 1995) and the RseA-family protein from P324 shares 67.63% amino acid identity with MucA from *P. aeruginosa* PAO1 and 34.54% amino acid identity to RseA from *E. coli*. In *P. aeruginosa*, mutations in *mucA* lead to increased transcription of *algR* and upregulation of alginate biosynthetic genes (Martin et al. 1993), which is the opposite effect of the *algR* mutation seen in P711. Furthermore, RpoE regulates biofilm formation, T6SS activity, and replication inside of macrophages in fish pathogen *P. plecoglossicida* (Zhang et al. 2023) and biofilm formation, stress tolerance, and survival within macrophages in human pathogen *Burkholderia pseudomallei* (Korbsrisate et al. 2005). Inactivation of RseA might therefore be expected to increase resistance to predation, but unregulated AlgU activity is toxic in *P. aeruginosa mucA* deletion mutants (Schofield et al. 2021) and *mucA* mutations reduce expression of many virulence genes (Rau et al. 2010). Altogether this suggests that biofilm formation may be more important for resistance in P711, which has a mucoid colony morphology, while other virulence factors are more important in P324.

### Reduced growth rate does not explain increased susceptibility to predation

While many of the genes we identified are likely to be involved in defense, some may have less direct effects. For example, dense bacterial lawns with a higher ratio of bacteria to amoebae are more resistant to predation (Adiba et al. 2010; DiSalvo et al. 2014), suggesting that mutations that reduce growth rate may increase susceptibility. To determine whether any of our predation-susceptible transposon mutants had growth defects, we measured growth curves and calculated the minimum doubling time for WT strains, all 6 mutants in P324, and 36 randomly selected P711 mutants (Figure 6). Among these strains, only 2 of the P324 mutants and none of the P711 mutants had doubling times that were significantly higher than WT, suggesting that loss of predation resistance is likely not due to reduced growth rate for most of the transposon mutants.

**Figure 6.**
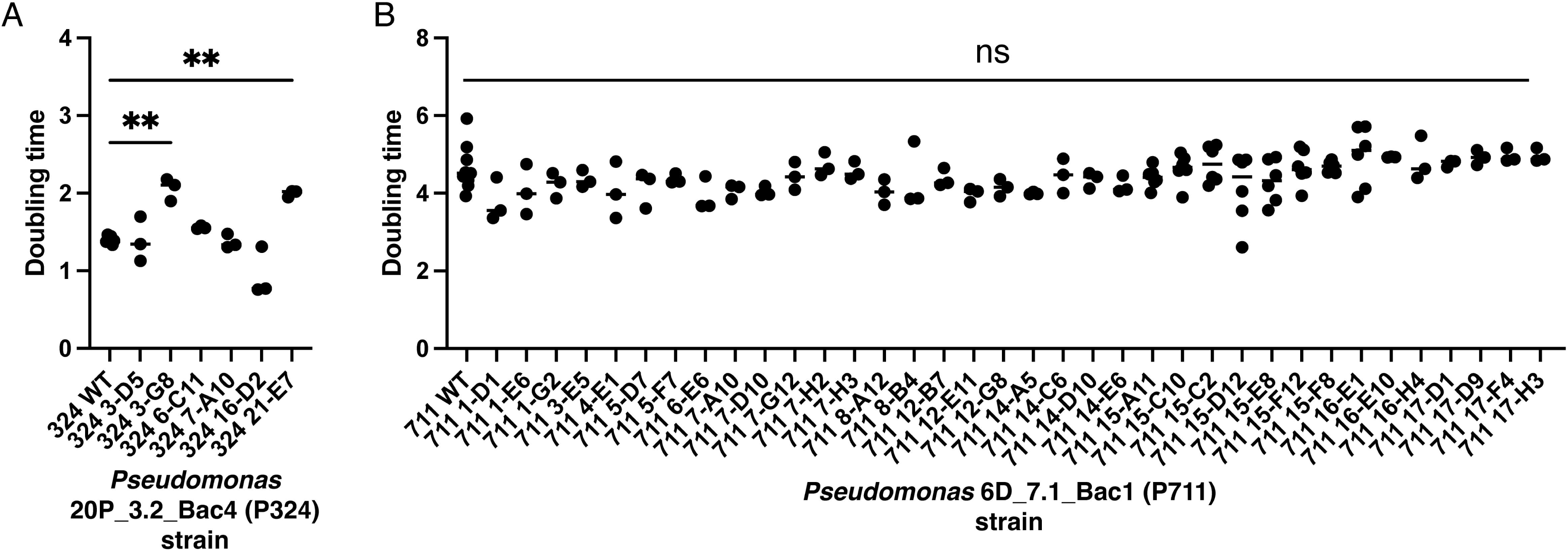
Minimum doubling times for wild type and transposon mutant strains. (A) *Pseudomonas* sp. P324 and (B) *Pseudomonas* sp. P711 WT and transposon mutant strains were grown in SM/5 broth and OD600 was measured every hour. Growthcurver was used to estimate the minimum doubling time for each strain. One-way ANOVA with Dunnett’s multiple comparisons test, p < 0.05.

## General Discussion

*Pseudomonas* sp. P324, *P. protegens* P821, and *Pseudomonas* sp. P711 are all soil bacteria that are resistant to predation by amoebae and capable of sporadically infecting *D. discoideum* fruiting bodies (Steele et al. 2023). The genome phylogeny for these species suggests that predation resistance has either evolved multiple times or been lost in many lineages. After screening more than 2,000 transposon mutants in each species, we identified few P324 and no P821 mutants that were susceptible to predation, suggesting that these strains have redundant mechanisms for fending off predators so that disruption of one gene is unlikely to compromise their resistance. This is supported by the observation that the few P324 mutants that were susceptible to predation had transposon insertions in genes with pleiotropic effects such as *gacA,* which regulates hundreds of genes including secondary metabolite biosynthetic pathways (Hassan et al. 2010), *dsbA*, which is required for correct folding of many secreted and periplasmic proteins, and *rseA/mucA*, which regulates the cell envelope stress response. In contrast, genes contributing to predation resistance in P711 are diverse in function but include multiple secondary metabolite biosynthetic gene clusters, toxins, a variety of transporters and transcriptional regulators, and genes predicted to have social functions.

It is interesting that the GacA-GacS two component system was required for predation resistance in all three species, given that the three species examined in this study appear to share no secondary metabolite biosynthetic gene clusters that are not also found in susceptible isolates (Steele et al. 2023). It seems that secondary metabolism overall is important for resistance, but each species may utilize different metabolites. Alternatively, secondary metabolism is not the only GacA regulated process that is important for resistance. Other predation resistance genes identified in P711 that are homologous to genes regulated by GacA in other *Pseudomonas* species include exoprotease AprA, alginate biosynthesis, and T6SS effectors (Lalaouna et al. 2012).

The importance of secreted metabolites and proteins for predation resistance is consistent with previous observations that resistant *Pseudomonas* species can protect otherwise susceptible bacteria from consumption by *D. discoideum* (Steele et al. 2023). Although the predation resistance genes we identified are not more likely to encode secreted or periplasmic proteins than we would expect based on the frequency of such genes in the genome, resistance genes in P711 were enriched in genes predicted to have social functions. We also observed that wild type P324 could rescue some susceptible mutants from predation, indicating that resistance is at least partially a public good. Furthermore, *D. discoideum* fruiting bodies infected by P324 are prone to secondary infections by prey bacteria, which would otherwise be consumed by the amoebae (Steele et al. 2023). Predation resistant Gac+ and susceptible Gac-*P. protegens* strains were co-isolated from naturally infected *D. discoideum*, suggesting Gac+ strains protect and facilitate the transmission of Gac-strains in nature (Stallforth et al. 2013). One of the assumptions of the Black Queen Hypothesis, which explains reductive genome evolution in free-living bacteria, is that behaviors bacteria engage in to protect themselves can be leaky and unintentionally benefit other species (Morris et al. 2012). One possible explanation for the distribution of predation resistance and related genes across the *Pseudomonas* phylogeny is that some species are exempted from selective pressure to maintain metabolically expensive mechanisms of resistance due to the absence of predators or the presence of resistant bacteria in the community. This could lead them to maintain regulatory pathways while losing downstream predation resistance genes such as toxins and secondary metabolite biosynthetic genes. However, keeping the regulatory machinery makes loss of predation resistance reversible through horizontal acquisition of new resistance genes (Costa et al. 2009; Lee et al. 2022). This could explain why some of the predation resistance genes in P711 have homologs in distantly related *Pseudomonas* species but not in close relatives.

Many genes required for *Pseudomonas* species to evade predation by amoebae, such as the GacA-GacS two-component system and the genes it regulates, also contribute to inhibition of fungal and bacterial competitors (Jousset et al. 2008; Lalaouna et al. 2012; Yan et al. 2018). These genes are required for the plant-beneficial activity *Pseudomonas* species used for biocontrol, but this activity is easily lost through spontaneous mutations in regulatory genes (Seaton et al. 2013; Stallforth et al. 2013). The presence of predators in microbial ecosystems therefore enforces the maintenance of genes relevant to human interests (Jousset et al. 2009; Rosenberg et al. 2009). It is not currently known whether secondary metabolites produced by our isolates are also effective against bacteria or fungi. However, the importance of GacA for resistance suggests that predation is likely to select against mutations that would inactivate this metabolically expensive regulon.

## Materials and Methods

Strains and plasmids used in this study are listed in Table 1. Antibiotic concentrations were 100 µg/mL carbenicillin (Crb), 300 µg/mL streptomycin (Str), 60 µg/mL spectinomycin (Sp), 20 µg/mL gentamicin (Gm), 30 µg/mL kanamycin (Km), 0.3 mM 4,6-diaminopimelic acid (DAP).

### Transposon mutagenesis

A mariner transposon that inserts into random sites in the genome was introduced into *Pseudomonas* strains P324, P821, and P711 through conjugation, using *Escherichia coli* β2163 + pBT20 as the donor, as previously described (Powell et al. 2016). Transformation efficiency was estimated by spotting serial dilutions of cells recovered from conjugations on LB and LB Gm and counting the colony forming units (CFU).

### Identification of predation-susceptible mutants

As we predicted that the mechanism of predation resistance was likely to be at least partially attributable to secreted molecules, we tested individual transposon mutants for loss of predation resistance. Pooled transposon mutant libraries were diluted in KK2 (2.25 g KH_2_PO_4_ (Sigma-Aldrich) and 0.67 g K_2_HPO_4_ (Fisher Scientific) per liter) and spread on LB Gm. Single colonies were collected and used to inoculate 100µl LB Gm broth in wells of 96-well plates, which were then incubated at 30°C overnight. *D. discoideum* AX4 amoebae were grown axenically in HL5 including glucose (Formedium) with Crb and Str. Cultures were centrifuged at 300 rcf for 5 min at 10°C, washed twice with 10 ml cold KK2, then resuspended in KK2 to a concentration of 8×10^6^ amoebae/ml. To identify mutants with increased susceptibility to predation, 50µl of overnight culture of each transposon mutant was mixed with 10µl of amoebae. 3µl droplets of each mixture were spotted on SM/5 agar plates, which were incubated at room temperature and monitored daily for consumption of bacteria and formation of fruiting bodies. 96-well plates containing overnight cultures of individual transposon mutants were cryopreserved by mixing 50 µl of bacterial culture with 12.5 µl of 80% glycerol before plates were stored at -80°C.

Some *P. protegens* mutants formed rings when grown with AX4, suggesting the amoebae consumed the bacteria at the center of the spot. However, this phenotype was not consistently reproducible and AX4 never formed fruiting bodies, indicating the mutants remained toxic. The density of the bacteria relative to the amoebae can affect the outcome of predation, a phenomenon known as an Allee effect (DiSalvo et al. 2014). To determine whether *P. protegens* mutants were susceptible to predation when grown at lower densities, we diluted cultures of a subset of ring-forming mutants to 20% or 2% of their original concentration before mixing them with AX4 and spotting the mixtures on agar, as described above. We did not observe any evidence that the diluted mutants were more susceptible to predation.

### Semirandom PCR

Semirandom PCR and Sanger sequencing were used to identify the location of transposon insertions in predation-susceptible mutants, as previously described (Powell et al. 2016). Briefly, cells were diluted in 200 μl nuclease-free water, boiled for 10 min to release DNA, then centrifuged for 30 s at 12,000 rcf to remove cell debris. Primers ARB-1B (5’-ggccagcgagctaacgagacnnnngatat-3’) and rnd1TnM20 (5’-tataatgtgtggaattgtgagcgg-3’) were used to amplify DNA adjacent to the transposon. PCR was performed by heating reactions at 95°C for 3 min, followed by 30 cycles of 95°C for 15 s, 33°C for 30 s, and then 72°C for 1 min 30 s, ending with a 3 minute final extension at 72°C. PCR products were diluted by mixing 5 µl with 195 µl of nuclease-free water, then additional amplification was performed using primers inARB-1B (5’-ggccagcgagctaacgagac-3’) and rnd2TnM20 (5’-acaggaaacaggactctagagg-3’). Reactions were heated to 95°C for 3 minutes, followed by 30 cycles of 95°C for 30 seconds, 56°C for 30 seconds, and then 72°C for 1 minute 30 seconds, ending with 3 minutes at 72°C. PCR products were column purified and submitted for Sanger sequencing with primer TnMSeq (5’-cacccagctttcttgtacac-3’). When semirandom PCR produced more than one product, the most abundant product was purified by excising the brightest band from a 1% agarose gel and extracting DNA using the NEB Monarch DNA Gel Extraction Kit. Gel purified PCR products were ligated into plasmid pGEM-T using the pGEM-T easy kit (Promega) and the resulting plasmid was transformed into *Escherichia coli* DH5α through electroporation. Plasmids were purified from DH5α using the Invitrogen plasmid miniprep kit and then Sanger sequenced by Genewiz (Azenta) using primers M13F (5’-GTAAAACGACGGCCAG-3’) and M13R (5’-CAGGAAACAGCTATGAC-3’). Sequencing reads were mapped to the bacterial genomes using Geneious v10.2.6.

### Identification of homologs in other *Pseudomonas* genomes

Protein-protein BLAST v2.9.0 (Camacho et al. 2009) was used to search *Pseudomonas* reference genomes and *D. discoideum* isolates for homologs to predation resistance genes from *Pseudomonas* P324 and P711. R v4.3.1 (R Core Team 2014) was used to construct a heatmap showing the amino acid percent identity of the best hits with >70% coverage. Orthofinder v2.5.4 (Emms and Kelly 2015; Emms and Kelly 2017; Emms and Kelly 2019) was used to construct a strain phylogeny based on 1703 gene trees.

### Phylogenies and tanglegrams

Genomes for the phylogeny were chosen to represent previously characterized *Pseudomonas* clades (Lalucat et al. 2020). Additional genomes were added by searching NCBI databases for homologs to the genes hypothesized to be horizontally transferred. We chose 78 conserved housekeeping genes from the Up-to-date Bacterial Core Gene set (UBCG) gene list (Na et al. 2018). Protein-protein BLAST and blastdbcmd v2.9.0 (Camacho et al. 2009) were used to extract sequences of homologous proteins, which were then aligned using MUSCLE (Edgar 2004). Alignments of core genes were concatenated using Geneious. RAxML (Stamatakis 2006; Stamatakis 2014) was used to construct maximum likelihood phylogenies from the protein alignments using the gamma model of rate heterogeneity and the LG amino acid substitution matrix with 1000 bootstraps. The resulting species tree was visualized using iToL (Letunic and Bork 2021). To investigate the role of horizontal gene transfer in acquisition of predation resistance genes, we chose four proteins that were found in few genomes that had annotations with clear connections to predation resistance. We used protein BLAST to identify homologs, MUSCLE to create alignments, and IQ-TREE (Nguyen et al. 2015; Minh et al. 2020) with model optimization (Kalyaanamoorthy et al. 2017) to build maximum likelihood phylogenies with 1000 bootstraps. Tanglegrams were visualized using dendextend v1.17.1 (Galili 2015) in R v4.4.0 (R Core Team 2014).

### Enrichment analysis

Clusters of Orthologous Genes (COG) functional categories were assigned to genes using eggNOG-mapper v2.1.12 (Huerta-Cepas et al. 2019; Cantalapiedra et al. 2021). pSORTb v 3.0 was used to predict the subcellular locations of proteins (Yu et al. 2010). SOCfinder v1.0 was used to identify cooperative genes (Belcher et al. 2023). To determine whether any categories were overrepresented among predation-resistance genes, Fisher’s exact test was used to compare the prevalence of predation-resistance genes assigned to each category to the prevalence of genes in that category in the genome as a whole.

### Targeted deletion of *gacA*

*Pseudomonas* P324 and *P. protegens* P821 Δ*gacA* mutants were created by replacing the *gacA* gene with a kanamycin resistance gene. Plasmids for homologous recombination were created using Golden Gate Assembly. DNA regions upstream and downstream of *gacA* were amplified using Phusion polymerase (NEB) and amplicons were purified using the GeneJET PCR Purification Kit (Thermo Scientific). 10 fmol of each PCR product and plasmids pYTK002, pYTK072, pBTK229, and pYTK095 were mixed with restriction enzyme BsaI-HFv2 (NEB), T4 DNA ligase (Promega), and 10X T4 DNA ligase buffer (Promega). Mixtures were incubated at 37°C for 1.5 min, then 16°C for 3 min for 30 cycles, then 37°C for 5 min and 80°C for 5 min. 2 µl of the Golden Gate Assembly reaction were electroporated into *E. coli* DH5α. Transformants were grown in LB supplemented with 100 µg/mL carbenicillin and 30 µg/mL kanamycin. Plasmid DNA was extracted using the NEB Monarch plasmid purification kit. 10 fmol of each plasmid was mixed with 10 fmol plasmid pCS1, restriction enzyme BsmBI-HF, T4 DNA ligase, and T4 DNA ligase buffer. Reactions were incubated at 42°C for 1.5 min and 16°C for 3 min for 30 cycles, then 50°C for 5 min and 80°C for 5 min. 2 µl of the reaction was electroporated into *E. coli* EC100D. Transformants were grown in LB supplemented with 60 µg/mL spectinomycin and 30 µg/mL kanamycin, then plasmid DNA was purified. PCR and Sanger sequencing were used to verify that plasmids were correctly assembled. Plasmids were then electroporated into *E. coli* MFD*pir*, a DAP auxotroph, and transformants were grown on LB supplemented with 60 µg/mL spectinomycin and 30 µg/mL kanamycin and 0.3 mM 2,6-diaminopimelic acid (DAP).

Conjugation was used to transform plasmids into *Pseudomonas* strains. Approximately 8×10^8^ donor and recipient cells were washed three times with KK2, then mixed and spotted on LB DAP plates. Conjugation plates were incubated at 30°C overnight, then cells were collected and washed three times with KK2, then spread on LB 30 µg/mL kanamycin plates. Transconjugants were streaked out on LB plates supplemented with 30 µg/mL kanamycin, 15 µg/mL tetracycline, and 1 µg/mL anhydrotetracycline, which induces expression of a toxin encoded by a gene on the pCS1 backbone used for counterselection. Replacement of *gacA* with the kanamycin resistance gene was verified through PCR.

### Fitness assays

Fitness assays were performed to compare the susceptibility of WT strains, Δ*gacA* deletion mutants, and transposon mutants to predation by *D. discoideum.* Bacteria were grown on LB agar, then collected and diluted to OD 1.5 in KK2. *D. discoideum* AX4 amoebae from axenic cultures were washed three times with cold KK2, then concentrated to 8×10^6^ cells/mL. For fitness assays with *D. discoideum* QS157, spores were collected from fruiting bodies and suspended in KK2 at a concentration of 2×10^6^ spores/mL. 65 µl of bacteria and 15 µl of *D. discoideum* were spread on 12 mL SM/5 plates. All plates were incubated at room temperature for 5 days. Plates were washed with 5 mL of KK2 to recover the remaining bacteria and serial dilutions of the recovered cells were spotted on LB agar. CFU were counted after 1-2 day incubation at 30°C.

### Rescue assays

Rescue assays were used to determine whether predation resistant wild type *Pseudomonas* could rescue susceptible transposon mutants from predation by *D. discoideum*. Kanamycin resistant, GFP-labeled wild type *Pseudomonas* P324 and gentamicin resistant transposon mutants were grown overnight at 30°C on LB plates. Cells were collected and suspended in KK2 at OD600 1.5, then mixed at a 50:50 ratio. 325µl of the bacterial suspension was mixed with 75µl of *D. discoideum* QS157 spores suspended in KK2 at a concentration of 2×10^6^ spores/mL. 80µl of bacteria and spores or 65µl of bacteria without spores were spread on 60mm SM/5 plates. Cells were collected after 5 d and serial dilutions were prepared and spotted on LB Km and LB Gm to enumerate the wild type and mutant *Pseudomonas* CFU remaining.

### Growth curves

Bacteria were collected from LB agar plates then diluted to OD 0.1 in SM/5 broth. 200 µl volumes were placed in a 96-well plate in triplicate. A Tecan Spark plate reader was used to measure the OD of each well every hour for 48 h. Growthcurver v0.3.1 (Sprouffske and Wagner 2016) was used to calculate the minimum doubling time of each strain.

## Acknowledgements

We thank Solange Kafunda for assistance with semirandom PCR, Emily Oxender for assistance with fitness assays, Israt Jahan for installing SOCfinder, and members of the Queller/Strassmann lab for useful discussions. We thank the Dicty Stock Center, Jeffery Barrick, Andrew Goodman, and John Dueber for providing strains and plasmids.

This manuscript is based upon work supported by the National Science Foundation (DEB 1753743 and DEB2237266 to DCQ and JES and NSF Postdoctoral Research Fellowships in Biology Program Grant No. 2109487 to MIS).

## Supplementary Figures and Tables

**Table S1. Genes disrupted by transposon insertions in predation-susceptible mutants.**

**Figure S1. Distribution of four predation resistance genes across *Pseudomonas* species.** The species phylogeny was constructed from an alignment of 78 conserved proteins from the Up-to-date Bacterial Core Gene set (UBCG) gene list. A maximum likelihood phylogeny was constructed using the gamma model of rate heterogeneity and the LG amino acid substitution matrix with 1000 bootstraps. Nodes with less than 90% bootstrap support are labeled, unlabeled nodes have >90% bootstrap support. Symbols on the right side of the phylogeny indicate the presence and absence of four predation resistance genes in each genome. Red: a metallohydrolase predicted to be a social gene. Orange: a predicted lysosome-like Type VI secretion system (T6SS) effector. Green: a gene predicted to play a role in secondary metabolism, located within a non-ribosomal peptide synthetase (NRPS) cluster. Blue: a predicted toxin with a YopT-type cysteine protease domain and a TcdA/TcdB pore forming domain.

**Figure S2. Phylogenies of four predation resistance genes are not congruent with the species phylogeny.** The species phylogeny is based on concatenated alignments of 78 core proteins from the UBCG gene list. Maximum likelihood phylogenies were constructed using IQ-TREE with model optimization and 1000 bootstraps. Tanglegrams were used to compare phylogenies for individual predation resistance proteins to the species tree. (A) A metallohydrolase predicted to be a social gene. (B) A predicted lysosome-like Type VI secretion system (T6SS) effector. (C) A gene predicted to play a role in secondary metabolism, located within a non-ribosomal peptide synthetase (NRPS) cluster. (D) A predicted toxin with a YopT-type cysteine protease domain and a TcdA/TcdB pore forming domain.

**Figure S3. *gacA* mutants reach higher abundance than WT on plates without *D. discoideum* AX4.** Strains were grown on SM/5 medium for 5 d. CFU were quantified by spotting a 1:10 dilution series on LB agar. P324: P324 6-C11, transposon insertion in *gacA*. P711: P711 8-G5, transposon insertion in *gacA*; P711 1-E6, NRPS-like cluster; P711 3-E5, NRPS/betalactone cluster; P711 8-H4, hserlactone/butyolactone cluster; P711 14-C6, NRPS/betalactone cluster; P711 16-E1, T1PKS cluster; P711 17-D1, NRPS-like cluster; P711 10-D3, *aprA*.

**Figure S4. *Pseudomonas* CFU recovered from plates with *D. discoideum* QS157.** Strains were grown on SM/5 medium with *D. discoideum* QS157 spores and bacteria were recovered after 5 days and enumerated by plating serial dilutions. QS157 is a wild clone that lacks laboratory adaptations seen in AX4. (A) *Pseudomonas* P324, (B) *P. protegens* P821, and (C) *Pseudomonas* P711 WT (red) and mutants lacking either *gacA* (orange), secondary metabolism biosynthetic gene clusters (green), or exoprotease *aprA* (blue) grown with *D. discoideum* QS157. One-way ANOVA with Tukey’s multiple comparisons test. Different letters indicate P value < 0.05.

