## Supplemental Table 1 for "The genetic basis of predation resistance in *Pseudomonas* species associated with the bactivorous soil amoeba *Dictyostelium discoideum*"

| **Supplementary Table 1. Genes disrupted by transposon insertions in predation-susceptible mutants.** | **Secondary metabolism (antiSMASH)** | No | NRPS-like cluster | No | No | No | No | No | No | No | No | No | No |
| --- | --- | --- | --- | --- | --- | --- | --- | --- | --- | --- | --- | --- | --- |
|  | **Protein Localization (pSORTb)** | Cytoplasmic | Cytoplasmic | Unknown | Extracellular | Cytoplasmic Membrane | Cytoplasmic | Cytoplasmic Membrane | Cytoplasmic | Cytoplasmic Membrane | Cytoplasmic | Unknown. Signal peptide detected | Cytoplasmic |
|  | **Social (SOCfinder)** | No | No | No | kofam: serralysin [EC:3.4.24.40] | No | No | No | No | No | No | No | No |
|  | **COG category** | S | S | K | Q | C | O | T | - | CO | T | H | S |
|  | **% of 33 strains with homologs** | 66.7 | 60.6 | 87.9 | 63.6 | 87.9 | 100.0 | 100.0 | 21.2 | 100.0 | 97.0 | 100.0 | 100.0 |
|  | **Category** | Toxin | Secondary metabolism | Transcriptional regulation | Toxin | Metabolism | Cellular processes and signaling | Transcriptional regulation | Toxin | Cellular processes and signaling | Transcriptional regulation | Metabolism | Uncharacterized |
|  | **Description** | Bacteriocin |  | Xenobiotic response element family of transcriptional regulators | Serralysin family metalloprotease | Cytochrome c oxidase polypeptide II (EC 1.9.3.1) | Belongs to the peptidase S16 family | Tricarboxylate transport sensor protein TctE. | T6SS effector cluster: Immediately upstream of Hcp and DUF2333 genes, upstream of VgrG and DUF 4123 gene | Protein-disulfide reductase DsbD | Two-component transcriptional response regulator, LuxR family. Part of chemotaxis receptor operon. | Catalyzes oxidation of 1,2-dihydro- and 1,6- dihydro- isomeric forms of beta-NAD(P) back to beta-NAD(P). | Protein of unknown function (DUF2797) |
|  | **Annotation** | Baseplate J/gp47 family protein | Diiron oxygenase | Helix-turn-helix domain-containing protein | *aprA* | *coxB* | AAA family ATPase | *tctE* | Hypothetical protein | *dsbD* | *pleD* | NAD(P)/FAD-dependent oxidoreductase | Hypothetical protein |
|  | **Locus tag** | NTD87_19285 | NTD87_08245 | NTD87_18900 | NTD87_17220 | NTD87_02005 | NTD87_15825 | NTD87_14210 | NTD87_18655 | NTD87_18675 | NTD87_18665 | NTD87_10290 | NTD87_22680 |
|  | **RefSeq Protein** | WP_003172142.1 | WP_003220441.1 | WP_054593225.1 | WP_262115219.1 | WP_262111506.1 | WP_262114939.1 | WP_262114578.1 | WP_262115579.1 | WP_262115584.1 | WP_262115580.1 | WP_262113698.1 | WP_262116559.1 |
|  | **Mutant ID** | 711 1-D1 | 711 1-E6 | 711 10-C2 | 711 10-D3 | 711 10-D8 711 22-G6 | 711 10-E2 | 711 10-G5 | 711 11-C10 | 711 11-H7 | 711 12-B7 | 711 12-E11 | 711 14-A5 |
|  | **Strain** | *Pseudomonas* 6D 7.1_Bac1 | *Pseudomonas* 6D 7.1_Bac1 | *Pseudomonas* 6D 7.1_Bac1 | *Pseudomonas* 6D 7.1_Bac1 | *Pseudomonas* 6D 7.1_Bac1 | *Pseudomonas* 6D 7.1_Bac1 | *Pseudomonas* 6D 7.1_Bac1 | *Pseudomonas* 6D 7.1_Bac1 | *Pseudomonas* 6D 7.1_Bac1 | *Pseudomonas* 6D 7.1_Bac1 | *Pseudomonas* 6D 7.1_Bac1 | *Pseudomonas* 6D 7.1_Bac1 |

| **Supplementary Table 1. Genes disrupted by transposon insertions in predation-susceptible mutants, continued.** | **Secondary metabolism (antiSMASH)** | NRPS/ betalactone protocluster | No | No | No | No | No | No | No | No | RiPP-like | T1PKS | No |
| --- | --- | --- | --- | --- | --- | --- | --- | --- | --- | --- | --- | --- | --- |
|  | **Protein Localization (pSORTb)** | Cytoplasmic Membrane | Cytoplasmic Membrane | Cytoplasmic | Cytoplasmic | Unknown. Signal peptide detected | Periplasmic | Unknown | Cytoplasmic Membrane | Cytoplasmic Membrane | Cytoplasmic | Cytoplasmic Membrane | Unknown |
|  | **Social (SOCfinder)** | A_SOCK: NRP-metallophore | No | No | No | No | No | No | No | No | No | No | No |
|  | **COG category** | M | P | IQ | O | I | E | S | H | I | S | M | S |
|  | **% of 33 strains with homologs** | 100.0 | 90.9 | 100.0 | 97.0 | 100.0 | 100.0 | 27.3 | 39.4 | 36.4 | 84.8 | 3.0 | 75.8 |
|  | **Category** | Transporter | Transporter | Metabolism | Cellular processes and signaling | Metabolism | Transporter | Uncharacterized | Metabolism | Metabolism | Secondary metabolism | Secondary metabolism | Uncharacterized |
|  | **Description** | Belongs to the membrane fusion protein (MFP) (TC 8.A.1) family | Belongs to the monovalent cation proton antiporter 2 (CPA2) transporter (TC 2.A.37) family | Long-chain fatty acid--CoA ligase | Xanthine and CO dehydrogenases maturation factor, XdhC/CoxF family CDS |  | Polyamine ABC transporter substrate-binding protein | Cupin 2, conserved barrel domain protein | NAD(P)H-hydrate dehydratase | Similar to dihydrorhizobitoxine fatty acid desaturase (RtxC) | Belongs to the UPF0276 family | COG1368 Phosphoglycerol transferase and related proteins, alkaline phosphatase superfamily | DUF3313 domain-containing protein |
|  | **Annotation** | Efflux RND transporter periplasmic adaptor subunit | *kefB* | *alkK* | *xdhC* | START domain-containing protein | *potF* | DHCW motif cupin fold protein | *nnrD* | Fatty acid desaturase family protein | DUF692 domain-containing protein | Lipoteichoic acid synthase family protein | *ydcL* |
|  | **Locus tag** | NTD87_17440 | NTD87_12465 | NTD87_10465 | NTD87_18855 | NTD87_04440 | NTD87_01300 | NTD87_22895 | NTD87_14810 | NTD87_24190 | NTD87_13675 | NTD87_12560 | NTD87_23560 |
|  | **RefSeq Protein** | WP_262115269.1 | WP_262114237.1 | WP_262113755.1 | WP_262115656.1 | WP_262112156.1 | WP_262111315.1 | WP_262116473.1 | WP_262114715.1 | WP_262116707.1 | WP_262114472.1 | WP_262114259.1 | WP_262116600.1 |
|  | **Mutant ID** | 711 14-C6 | 711 14-D10 | 711 14-E6 | 711 15-A11 | 711 15-C2 | 711 15-E8 711 22-F8 | 711 15-F12 | 711 15-F8 | 711 16-B5 | 711 16-C9 | 711 16-E1 | 711 16-F9 |
|  | **Strain** | *Pseudomonas* 6D 7.1_Bac1 | *Pseudomonas* 6D 7.1_Bac1 | *Pseudomonas* 6D 7.1_Bac1 | *Pseudomonas* 6D 7.1_Bac1 | *Pseudomonas* 6D 7.1_Bac1 | *Pseudomonas* 6D 7.1_Bac1 | *Pseudomonas* 6D 7.1_Bac1 | *Pseudomonas* 6D 7.1_Bac1 | *Pseudomonas* 6D 7.1_Bac1 | *Pseudomonas* 6D 7.1_Bac1 | *Pseudomonas* 6D 7.1_Bac1 | *Pseudomonas* 6D 7.1_Bac1 |

| **Supplementary Table 1. Genes disrupted by transposon insertions in predation-susceptible mutants, continued.** | **Secondary metabolism (antiSMASH)** | NRPS-like | No | No | No | No | No | No | No | No | No | No | No |
| --- | --- | --- | --- | --- | --- | --- | --- | --- | --- | --- | --- | --- | --- |
|  | **Protein Localization (pSORTb)** | Unknown | Outer Membrane | Unknown | Cytoplasmic Membrane | Unknown | Cytoplasmic | Cytoplasmic | Unknown | Cytoplasmic Membrane | Cytoplasmic | Unknown | Cytoplasmic |
|  | **Social (SOCfinder)** | kofam: adenylylsulfate reductase, subunit B [EC:1.8.99.2] | Social | No | kofam: insecticidal toxin | No | No | No | No | No | No | No | No |
|  | **COG category** | U | M | E | Q | S | J | KL | O | CP | H | S | S |
|  | **% of 33 strains with homologs** | 24.2 | 100.0 | 100.0 | 12.1 | 3.0 | 100.0 | 3.0 | 18.2 | 100.0 | 100.0 | 3.0 | 100.0 |
|  | **Category** | Secondary metabolism | Cellular processes and signaling | Metabolism | Toxin | Uncharacterized | Information storage and processing | Information storage and processing | Cellular processes and signaling | Metabolism | Metabolism | Uncharacterized | Cellular processes and signaling |
|  | **Description** | Domain of unknown function (DUF4347) |  | Belongs to the peptidase S1B family | TcdA/TcdB pore forming domain |  | Class I SAM-dependent methyltransferase | DEAD-like helicase domains, catalytic domain of type II restriction enzymes | OsmC family protein | Catalyzes the transfer of electrons from NADH to ubiquinone | Catalyzes the oxidation of L-aspartate to iminoaspartate | Caudovirales tail fibre assembly protein, lambda gpK | Pyrimidine/purine nucleotide 5'-monophosphate nucleosidase PpnN (EC 3.2.2.4) (EC 3.2.2.10). |
|  | **Annotation** | Ig-like domain-containing protein | DUF1302 domain-containing protein | Serine protease | YopT-type cysteine protease domain-containing protein | FAD/NAD(P)-binding protein | *rlmG* | SNF2-related protein | *yhfA* | *nuoL* | *nadB* | Phage tail protein | *ppnN* |
|  | **Locus tag** | NTD87_08305 | NTD87_10470 | NTD87_05630 | NTD87_18740 | NTD87_20415 | NTD87_15915 | NTD87_17790 | NTD87_06935 | NTD87_27380 | NTD87_14235 | NTD87_22065 | NTD87_25035 |
|  | **RefSeq Protein** | WP_262113186.1 | WP_262113757.1 | WP_262112472.1 | WP_262115603.1 | WP_262115987.1 | WP_262114962.1 | WP_262115355.1 | WP_056857866.1 | WP_262117288.1 | WP_262114582.1 | WP_262116294.1 | WP_262116866.1 |
|  | **Mutant ID** | 711 17-D1 | 711 17-D9 | 711 17-F2 | 711 17-F4 | 711 17-F7 | 711 17-H3 | 711 18-B3 | 711 18-B6 | 711 18-C10 | 711 18-E8 711 18-E9 | 711 18-F10 | 711 19-A4 |
|  | **Strain** | *Pseudomonas* 6D 7.1_Bac1 | *Pseudomonas* 6D 7.1_Bac1 | *Pseudomonas* 6D 7.1_Bac1 | *Pseudomonas* 6D 7.1_Bac1 | *Pseudomonas* 6D 7.1_Bac1 | *Pseudomonas* 6D 7.1_Bac1 | *Pseudomonas* 6D 7.1_Bac1 | *Pseudomonas* 6D 7.1_Bac1 | *Pseudomonas* 6D 7.1_Bac1 | *Pseudomonas* 6D 7.1_Bac1 | *Pseudomonas* 6D 7.1_Bac1 | *Pseudomonas* 6D 7.1_Bac1 |

| **Supplementary Table 1. Genes disrupted by transposon insertions in predation-susceptible mutants, continued.** | **Secondary metabolism (antiSMASH)** | No | No | No |  | No | No | No | No | No | No | No | No |
| --- | --- | --- | --- | --- | --- | --- | --- | --- | --- | --- | --- | --- | --- |
|  | **Protein Localization (pSORTb)** | Cytoplasmic | Unknown | Cytoplasmic | Unknown | Cytoplasmic Membrane | Unknown | Cytoplasmic Membrane | Cytoplasmic Membrane | Cytoplasmic | Unknown . Signal peptide detected | Unknown. Signal peptide detected | Cytoplasmic |
|  | **Social (SOCfinder)** | No | No | No | No | No | No | No | kofam: undecaprenyl-phosphate glucose phosphotransferase [EC:2.7.8.31] | No | No | kofam: linear primary-alkylsulfatase [EC:3.1.6.21] | No |
|  | **COG category** | D | S | I | - | M | S | G | M | L | ET | Q | - |
|  | **% of 33 strains with homologs** | 15.2 | 100.0 | 48.5 | 33.3 | 97.0 | 81.8 | 100.0 | 72.7 | 100.0 | 93.9 | 18.2 | 3.0 |
|  | **Category** | Cellular processes and signaling | Uncharacterized | Metabolism | Uncharacterized | Transporter | Uncharacterized | Transporter | Cellular processes and signaling | Information storage and processing | Transporter | Social | Uncharacterized |
|  | **Description** | The Fic protein is involved in cell division |  | Serine aminopeptidase, S33 |  | BCCT family transporter | Amidohydrolase family | Major facilitator superfamily | Bacterial sugar transferase, capsular polysaccharide biosynthesis protein | Belongs to the DNA glycosylase MPG family | Amino acid ABC transporter substrate-binding protein | Alkyl sulfatase BDS1 and related hydrolases, metallo-beta-lactamase superfamily |  |
|  | **Annotation** | Fic family protein | LamB/YcsF family protein | Alpha/beta fold hydrolase | Hypothetical protein | *betT* | *nfdA* | *ampG* | *wcaJ* | DNA-3-methyladenine glycosylase | ABC transporter substrate-binding protein | MBL fold metallo-hydrolase | SLOG family protein |
|  | **Locus tag** | NTD87_14130 | NTD87_14530 | NTD87_03865 | NTD87_27790 | NTD87_07540 | NTD87_21485 | NTD87_13490 | NTD87_27105 | NTD87_16475 | NTD87_26555 | NTD87_19505 | NTD87_17775 |
|  | **RefSeq Protein** | WP_262114562.1 | WP_262114640.1 | WP_262112008.1 | WP_262117356.1 | WP_262113018.1 | WP_262116191.1 | WP_262114439.1 | WP_262117240.1 | WP_262115078.1 | WP_262117148.1 | WP_262115828.1 | WP_262115351.1 |
|  | **Mutant ID** | 711 19-A8 | 711 19-C4 | 711 19-D3 | 711 19-F5 | 711 19-G2 | 711 20-G9 | 711 21-E4 | 711 22-G9 | 711 5-F7 | 711 6-E6 | 711 7-A10 | 711 7-D10 |
|  | **Strain** | *Pseudomonas* 6D 7.1_Bac1 | *Pseudomonas* 6D 7.1_Bac1 | *Pseudomonas* 6D 7.1_Bac1 | *Pseudomonas* 6D 7.1_Bac1 | *Pseudomonas* 6D 7.1_Bac1 | *Pseudomonas* 6D 7.1_Bac1 | *Pseudomonas* 6D 7.1_Bac1 | *Pseudomonas* 6D 7.1_Bac1 | *Pseudomonas* 6D 7.1_Bac1 | *Pseudomonas* 6D 7.1_Bac1 | *Pseudomonas* 6D 7.1_Bac1 | *Pseudomonas* 6D 7.1_Bac1 |

| **Supplementary Table 1. Genes disrupted by transposon insertions in predation-susceptible mutants, continued.** | **Secondary metabolism (antiSMASH)** | No | redox-cofactor protocluster | No | No | No | No | No | No | No | No | No | No |
| --- | --- | --- | --- | --- | --- | --- | --- | --- | --- | --- | --- | --- | --- |
|  | **Protein Localization (pSORTb)** | Unknown | Cytoplasmic | Cytoplasmic Membrane | Cytoplasmic Membrane | Cytoplasmic Membrane | Cytoplasmic | Cytoplasmic | Unknown Signal peptide detected | Periplasmic | Unknown Signal peptide detected | Cytoplasmic Membrane | Cytoplasmic |
|  | **Social (SOCfinder)** | No | No | No | No | No | kofam: two-component system, response regulator AauR | No | No | No | No | No | No |
|  | **COG category** | M | K | S | P | E | T | K | P | ET | C | EGP | C |
|  | **% of 33 strains with homologs** | 18.2 | 100.0 | 97.0 | 90.9 | 100.0 | 100.0 | 100.0 | 60.6 | 100.0 | 93.9 | 100.0 | 97.0 |
|  | **Category** | Toxin | Transcriptional regulation/secondary metabolism | Uncharacterized | Transporter | Transporter | Transcriptional regulation | Transcriptional regulation | Metabolism | Transporter | Metabolism | Transporter | Metabolism |
|  | **Description** | Hypothetical gene downstream of VgrG, lysozyme-like domain. T6SS effector cluster. |  |  | OFA family MFS transporter | *Transposon is 4bp upstream of the gene | Fis family transcriptional regulator, two component regualator of acidic amino acid uptake. | LytTR family DNA-binding domain-containing protein | Alpha-2-macroglobulin family protein, large extracellular alpha-helical protein | Belongs to the bacterial solute-binding protein 3 family | cytochrome c | Facilitates transport across cytoplasmic or internal membranes of one or more from a variety of substrates | FAD-binding oxidoreductase |
|  | **Annotation** | Lysozyme | LysR family transcriptional regulator | TM2 domain-containing protein | *yhjX* | Phenylalanine-specific permease/amino acid permease | Sigma-54 dependent transcriptional regulator | *algR* | *yfhM* | Transporter substrate-binding domain-containing protein | *adhB* | MFS transporter | *lldE* |
|  | **Locus tag** | NTD87_07950 | NTD87_07210 | NTD87_24495 | NTD87_05595 | NTD87_04060 | NTD87_08990 | NTD87_09840 | NTD87_18605 | NTD87_00585 | NTD87_01975 | NTD87_10035 | NTD87_03665 |
|  | **RefSeq Protein** | WP_262113099.1 | WP_262112891.1 | WP_262116770.1 | WP_262112464.1 | WP_262112057.1 | WP_262113353.1 | WP_262113571.1 | WP_262115565.1 | WP_262111137.1 | WP_262111497.1 | WP_262113620.1 | WP_262111953.1 |
|  | **Mutant ID** | 711 7-G12 711 15-C10 | 711 7-H2 | 711 7-H3 | 711 8-A12 | 711 8-A4 711 16-B8 | 711 8-A8 | 711 8-A9 | 711 8-B4 | 711 8-C2 | 711 8-D10 | 711 8-D8 | 711 8-E2 |
|  | **Strain** | *Pseudomonas* 6D 7.1_Bac1 | *Pseudomonas* 6D 7.1_Bac1 | *Pseudomonas* 6D 7.1_Bac1 | *Pseudomonas* 6D 7.1_Bac1 | *Pseudomonas* 6D 7.1_Bac1 | *Pseudomonas* 6D 7.1_Bac1 | *Pseudomonas* 6D 7.1_Bac1 | *Pseudomonas* 6D 7.1_Bac1 | *Pseudomonas* 6D 7.1_Bac1 | *Pseudomonas* 6D 7.1_Bac1 | *Pseudomonas* 6D 7.1_Bac1 | *Pseudomonas* 6D 7.1_Bac1 |

| **Supplementary Table 1. Genes disrupted by transposon insertions in predation-susceptible mutants, continued.** | **Secondary metabolism (antiSMASH)** | No | No | gacA | No | No | hserlactone/ butytolactone cluster | No | No | No | No | No | No |
| --- | --- | --- | --- | --- | --- | --- | --- | --- | --- | --- | --- | --- | --- |
|  | **Protein Localization (pSORTb)** | Cytoplasmic | Periplasmic | Cytoplasmic | Unknown | Cytoplasmic | Cytoplasmic | Cytoplasmic Membrane | Cytoplasmic | Cytoplasmic Membrane | Cytoplasmic | Cytoplasmic | Periplasmic |
|  | **Social (SOCfinder)** | No | No | No | No | No | No | No | No | No | No | No | No |
|  | **COG category** | not found | ET | K | L | S | I | P | K | EG | K | G | O |
|  | **% of 33 strains with homologs** | 100.0 | 100.0 | 100.0 | 18.2 | 84.8 | 100.0 | 100.0 | 100.0 | 93.9 | 100.0 | 100.0 | 100.0 |
|  | **Category** | Transcriptional regulation | Transporter | Transcriptional regulation | Information storage and processing | Uncharacterized | Metabolism | Transporter | Transcriptional regulation | Transporter | Information storage and processing | Metabolism | Cellular processes and signaling |
|  | **Description** |  | Bacterial periplasmic substrate-binding proteins | Bacterial regulatory proteins, luxR family, gacA | Belongs to the 'phage' integrase family | Plant-induced nitrilase (EC 3.5.5.1) | acetyl-CoA C-acetyltransferase | ABC-type spermidine/putrescine transport system, permease component II | Putative choline sulfate-utilization transcription factor | carboxylate/amino acid/amine transporter | Transcriptional regulator, PLP-dependent aminotransferase family protein | Xylulose kinase, FGGY-family carbohydrate kinase | Thiol disulfide interchange protein |
|  | **Annotation** | Winged helix-turn-helix domain-containing protein | ABC transporter substrate-binding protein | *gacA/uvrY* | Hypothetical protein | Carbon-nitrogen hydrolase family protein | *fadI* | ABC transporter permease | LysR family transcriptional regulator | *yigM* | *yjiR* | *ygcE* | *dsbA* |
|  | **Locus tag** | NTD87_27120 | NTD87_18250 | NTD87_21580 | NTD87_17830 | NTD87_00150 | NTD87_18450 | NTD87_17635 | NTD87_01870 | NTD87_14695 | NTD87_10030 | NTD87_03935 | NTG44_27720 |
|  | **RefSeq Protein** | WP_262117243.1 | WP_262115470.1 | WP_019690936.1 | WP_262115363.1 | WP_262111040.1 | WP_262115525.1 | WP_262115316.1 | WP_262111464.1 | WP_262114674.1 | WP_262113619.1 | WP_262112026.1 | WP_095159042.1 |
|  | **Mutant ID** | 711 8-E5 | 711 8-G4 | 711 8-G5 | 711 8-G7 | 711 8-H3 | 711 8-H4 | 711 9-A8 | 711 9-C1 | 711 9-D6 | 711 9-D9 | 711 9-E5 | 324 1-B12 |
|  | **Strain** | *Pseudomonas* 6D 7.1_Bac1 | *Pseudomonas* 6D 7.1_Bac1 | *Pseudomonas* 6D 7.1_Bac1 | *Pseudomonas* 6D 7.1_Bac1 | *Pseudomonas* 6D 7.1_Bac1 | *Pseudomonas* 6D 7.1_Bac1 | *Pseudomonas* 6D 7.1_Bac1 | *Pseudomonas* 6D 7.1_Bac1 | *Pseudomonas* 6D 7.1_Bac1 | *Pseudomonas* 6D 7.1_Bac1 | *Pseudomonas* 6D 7.1_Bac1 | *Pseudomonas* 20P 3.2_Bac4 |

| **Supplementary Table 1. Genes disrupted by transposon insertions in predation-susceptible mutants, continued.** | **Secondary metabolism (antiSMASH)** | No | No | No | No |
| --- | --- | --- | --- | --- | --- |
|  | **Protein Localization (pSORTb)** | Cytoplasmic | Cytoplasmic | Cytoplasmic | Cytoplasmic Membrane |
|  | **Social (SOCfinder)** | No | No | No | No |
|  | **COG category** | C | ET | K | T |
|  | **% of 33 strains with homologs** | 100.0 | 93.9 | 100.0 | 100.0 |
|  | **Category** | Metabolism | Transporter | Transcriptional regulation | Transcriptional regulation |
|  | **Description** | Belongs to the citrate synthase family | Amino acid ABC transporter substrate-binding protein | Two component transcriptional regulator, LuxR family | Anti sigma-E protein RseA |
|  | **Annotation** | *prpC* | ABC transporter substrate-binding protein | *gacA/uvrY* | *rseA/mucA* |
|  | **Locus tag** | NTG44_09100 | NTG44_24410 | NTG44_12650 | NTG44_10810 |
|  | **RefSeq Protein** | WP_262154216.1 | WP_262156137.1 | WP_095159785.1 | WP_262154406.1 |
|  | **Mutant ID** | 324 16-D2 | 324 3-D5 | 324 6-C11 | 324 7-A10 324 21-E7 |
|  | **Strain** | *Pseudomonas* 20P 3.2_Bac4 | *Pseudomonas* 20P 3.2_Bac4 | *Pseudomonas* 20P 3.2_Bac4 | *Pseudomonas* 20P 3.2_Bac4 |
