## Supplementary figures and images for "The genetic basis of predation resistance in *Pseudomonas* species associated with the bactivorous soil amoeba *Dictyostelium discoideum*"

### Supplementary Figure 1

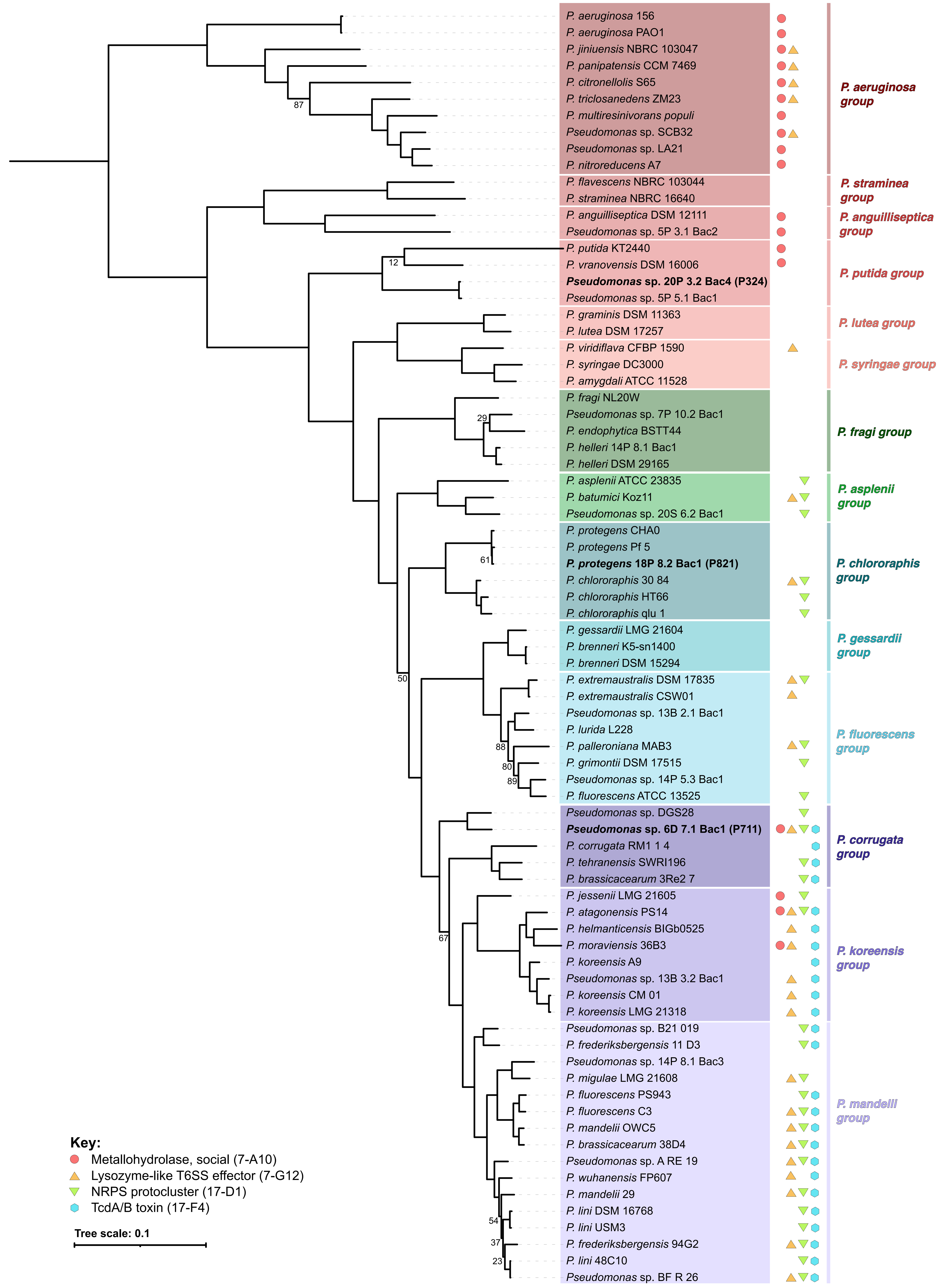

### Supplementary Figure 2

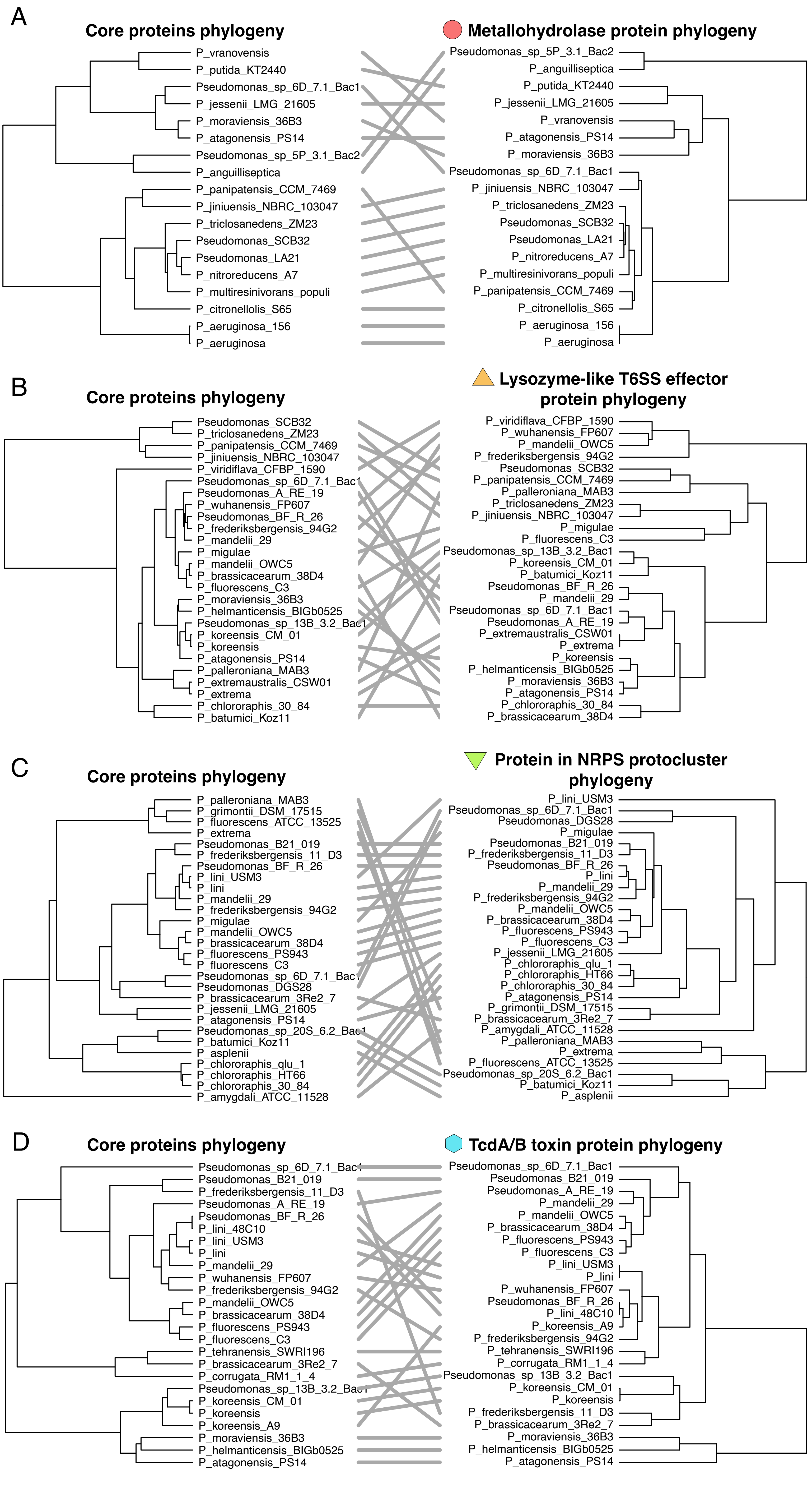

### Supplementary Figure 3

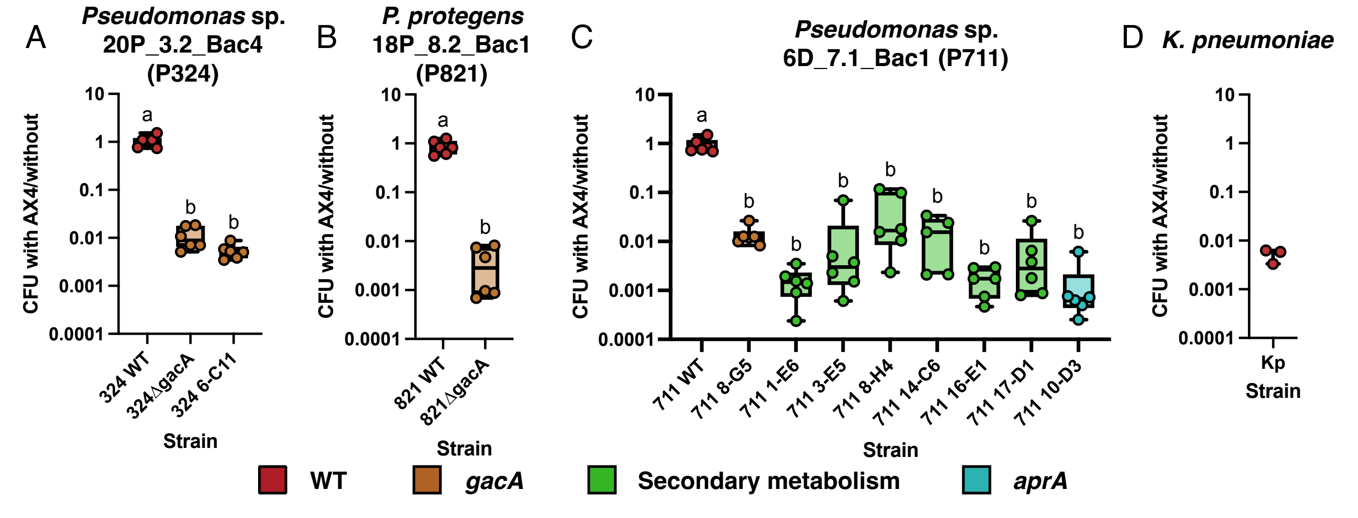

### Supplementary Figure 4

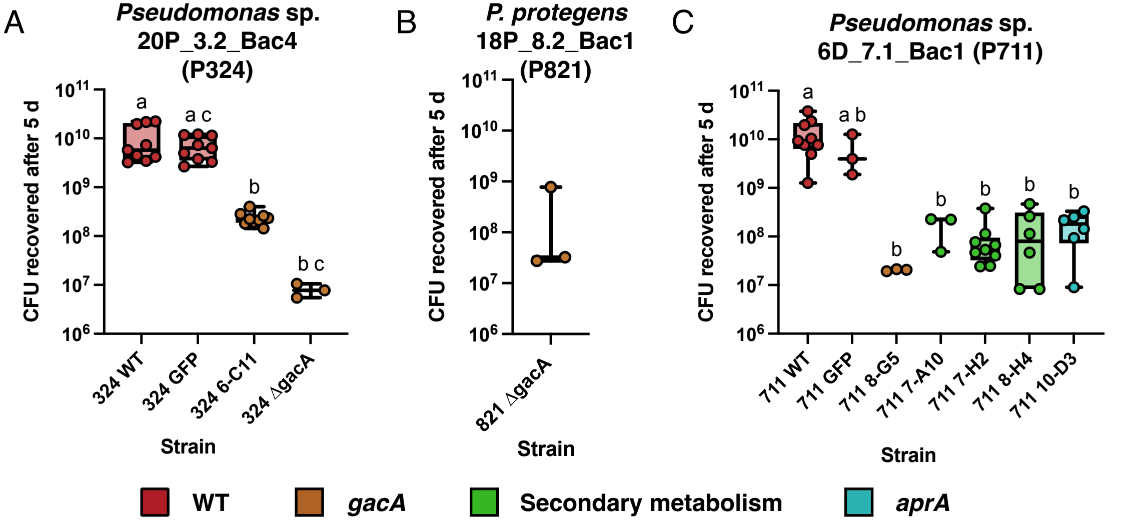
